# Neutrophil-chemoattractant CXCL5 induces lung barrier permeability in acute lung injury

**DOI:** 10.1101/2024.11.25.625215

**Authors:** Sarah Berger, Cengiz Goekeri, Peter Pennitz, Birgitt Gutbier, Laura Michalick, Karen Hoffmann, Elena Lopez-Rodriguez, Vladimir Gluhovic, Sandra-Maria Wienhold, Ulrike Behrendt, Alexander Taylor, Kristina Dietert, Holger Kirsten, Sandra Kunder, Kristina Mueller, Markus Weigel, Torsten Hain, Sarah M. Volkers, Sebastian Weis, Achim D. Gruber, Leif E. Sander, Wolfgang M. Kuebler, CAPSyS study group, Martin Witzenrath, Geraldine Nouailles

## Abstract

**Rationale:** Acute lung injury is frequently caused by pneumonia, and severity correlates with inflammatory cell recruitment and lung barrier failure. Deciphering drug-targetable pathways is of clinical importance.

**Objectives:** Investigation of paracrine and autocrine effects of epithelial-derived chemokine CXCL5 in acute lung injury.

**Methods:** We assessed the role of CXCL5 and related chemokines in patients with severe pneumonia and primary cells or cell lines challenged with *Streptococcus pneumoniae* or exposed to mechanical stretch. Furthermore, we evaluated the role of CXCL5 in *in vivo* models of acute lung injury through use of *Cxcl5*-deficient mice and *in vitro* human lung barrier models.

**Results:** Pneumococcal infection and mechanical ventilation were associated with CXCL5 production in human subjects and in mice. CXCL5 was produced by bronchial and alveolar epithelial cells. The alveolar-epithelial barrier was protected in acute lung injury models in *Cxcl5*-deficient mice, independent of alveolar neutrophil recruitment. Single-cell transcriptomics revealed enhanced cell junctional transcripts in epithelial, but not endothelial cells in *Cxcl5*-deficient mice. Accordingly, CXCL5 exposure disrupted the barrier function of TNF-primed human primary alveolar epithelial cells, but not pulmonary microvascular endothelial cells.

**Conclusions:** We describe a novel function of CXCL5 in acute lung injury. Besides its recognized role in recruiting highly inflammatory cells such as neutrophils, CXCL5 increases alveolar-epithelial barrier permeability. Thus, in severe bacterial pneumonia, targeting of CXCL5 as adjunctive therapy to antibiotics may aid in reducing overshooting inflammation as well as stabilizing lung barrier function.

## INTRODUCTION

Community-acquired pneumonia (CAP), frequently caused by *Streptococcus pneumoniae* (*S.pn.*), remains a considerable global health burden, with high incidence and mortality [1]. Severe CAP courses may result in acute lung injury (ALI) [2] and the need for mechanical ventilation for survival, with the risk of causing ventilator-induced lung injury (VILI) [3]. Hallmarks of ALI include excessive leukocyte influx, edema and alveolar-capillary barrier failure [4, 5]. Physiologically the lung barrier is maintained by alveolar epithelial cells and endothelial cells (EC) [6].

During pulmonary inflammation, neutrophils are rapidly recruited into alveolar spaces through chemo-attractants including, but not limited to CXCL-chemokines via G-protein coupled receptors such as CXCR2 [7, 8], whilst CXCL5 in particular is produced by lung epithelial cells to initiate neutrophil recruitment during inflammation and infection [9]. However, inflammatory cells, microbes or mechanical ventilation can disrupt lung barrier integrity, resulting in alveolar flooding [10]. Thus, despite the critical role of neutrophils in bacterial clearance, their effector functions remain a double-edged sword [11].

CXCR2 inhibitors have recently been trialed in a multicenter, phase II randomized controlled study in hospitalized patients with COVID-19 pneumonia (NCT04794803), revealing that treatment was well tolerated and improved the time to composite endpoint of clinical events (use of supplemental oxygen, need for mechanical ventilation, intensive care unit admission, and/or use of rescue medication) [12], while further trials in patients with ARDS (phase II clinical trial; NCT05496868) and severe CAP, including COVID-19 (phase III clinical trial; NCT05254990) are currently ongoing.

Previously, we demonstrated that targeted deletion of the gene encoding a CXCR2 ligand, the epithelial-derived *Cxcl5*, sufficed to limit detrimental neutrophil-mediated inflammation in pulmonary tuberculosis [13]. In pneumococcal pneumonia, CXCL5 can serve as a valid marker for epithelial cell activation [14], however its involvement in ALI remains lacking. Thus, we aimed to characterize the role of CXCL5 in regulating lung barrier permeability during pneumococcal pneumonia and VILI.

## MATERIALS AND METHODS

Additional detail on the Materials and Methods is provided in an Online Data Supplement.

### Ethics for patient samples

Data of severe CAP patients requiring mechanical ventilation were obtained from the CAPSyS observational study (sub-study of PROGRESS, clinicaltrials.gov: NCT02782013) [15], which was a non-interventional prospective study, as approved by the Ethics Committee of the University of Jena (2403-10/08) and the local Ethics Committee of each study site. All participants or their legal guardians provided written informed consent to participate in the study. The study met the requirements of the Declaration of Helsinki and ICH-GCP guidelines.

In the non-CAP control cohort, use of bronchoalveolar lavage fluid (BALF) samples for scientific purposes received approval and oversight from the Ethics Committee of the Charité - Universitätsmedizin Berlin, Germany (EA2/086/16).

### Ethics for animal experiments

Institutional and governmental (LaGeSo) authorities approved animal procedures. FELASA guidelines were followed. Female C57BL/6J (WT) and *Cxcl5*^−/−^ mice, aged 8-10 weeks, were housed under specific-pathogen-free conditions. Mice undergoing experimental procedures were monitored for body temperature and weight changes and scored for humane end-point criteria at ∼12h intervals. Mice reaching pre-defined end-point criteria were euthanized before developing severe symptoms.

### Murine pneumococcal pneumonia

For infections, mice anesthetized with ketamine and xylazine were transnasally inoculated with 5 × 10^6^ colony forming units (CFU) *S.pn.* serotype (ST)2 (NCTC 7466) or ST3 (NCTC 7978), while control mice received sham (1×PBS) [16]. Prior to dissection at 24 hours post-infection (hpi) mice were deeply anesthetized by ketamine and xylazine, checked for loss of intertoe reflexes and euthanized by exsanguination.

### Murine ventilator-induced lung injury (VILI)

Prior to ventilation, mice were anesthetized with fentanyl, midazolam, and medetomidine. Briefly, mice received high tidal volume (HVT) ventilation with a tidal volume of 34 ml/kg body weight, respiratory rate of 70/min for 4h. Control non-ventilated (NV) mice, were ventilated for 5-10 min at tidal volume of 9 ml/kg body weight, respiratory rate of 160/min for baseline measurements [17]. Mice were sacrificed though exsanguination via the carotic catheter. Human serum albumin (HSA) was applied ninety minutes before the termination of the experiment, as previously described [18].

### Pneumolysin treatment of isolated perfused mouse lungs (IPML)

Mice were intraperitoneally anesthetized with ketamine, xylazine admixed in a heparin saline solution (50% heparin) and then exsanguinated. Tracheotomy and ventilation, isolated perfusion, baseline generation and HSA application were performed as described [19]. Pneumolysin was either intratracheally aerosolized by MicroSprayer^®^ (Penn-Century, USA; 1 µg in 25 µl bolus) or intravenously administered by perfusion (1.75 µg in 1 ml bolus). Control groups received corresponding amount of buffer. Thirty minutes after challenge, BAL was performed, and the HSA concentration was measured in BALF via ELISA according to manufacturer’s instructions (Bethyl Laboratories, Montgomery, TX, USA).

## RESULTS

### Patient characteristics

6 patients with CAP and confirmed *S.pn.* infection – identified by nanopore 16S rRNA sequencing – were included from the CAPSyS cohort, whilst 5 non-CAP patients were included as controls. The clinical and baseline characteristics are shown in Table E1. BAL was performed on the same day (n=3), one day after (n=2) or two days after hospitalization (n=1) in patients with CAP. No antibiotic treatment prior to hospitalization was documented. BALF samples in non-CAP controls were obtained from individuals who had undergone bronchoscopy for the exclusion of pulmonary tuberculosis, pulmonary involvement in systemic disease or due to idiopathic coughing.

### Pneumococcal infection and mechanical ventilation induce epithelial-derived CXCL5 production

In the BALF of mechanically-ventilated *S.pn.*-infected CAP patients, hCXCL5 and hCXCL8 (IL-8) were significantly elevated compared to non-CAP patients (Figure 1A, Table E1). In mice, CXCL5 represented the highest chemokine concentration 24h and 48h following *S.pn.* infection (Figure 1B). Similarly, in a murine model of VILI, after 4 hours of ventilation, CXCL5 was the predominant chemokine detected in BALF (Figure 1C).

**Figure 1.**
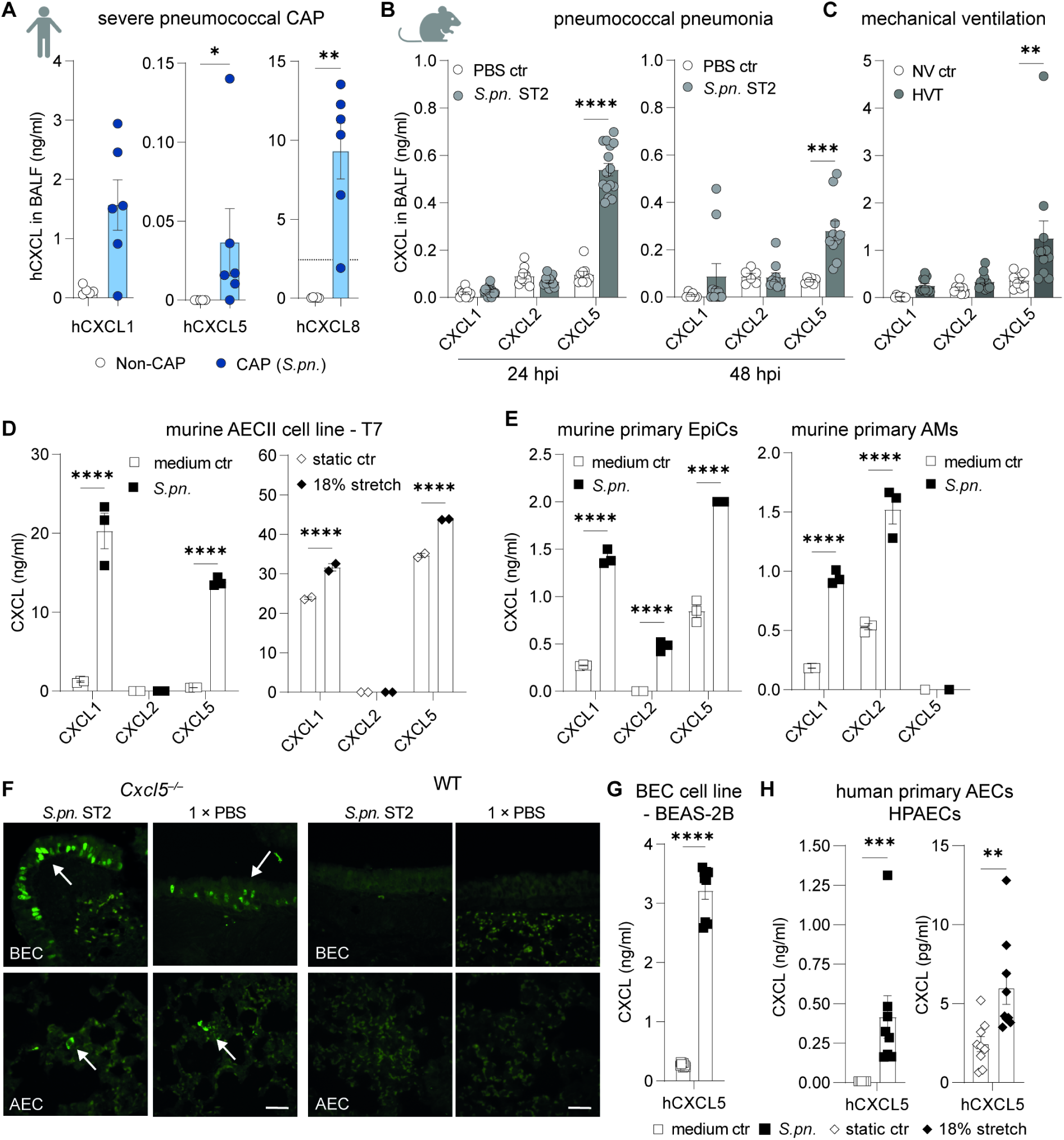
CXCL-chemokine production is induced in patients with severe pneumococcal CAP, lung epithelial cells and models of acute lung injury. CXCL-chemokine concentrations were measured by ELISA in BALF of (**A**) patients with severe pneumococcal CAP (n = 6) and non-CAP controls (n = 5), in addition to (**B**) mice intranasally infected with *S. pneumoniae* serotype 2 (*S.pn.* ST2) or treated with PBS (PBS ctr), which were sacrificed at 24- (n_PBS_ = 9, n*_S.pn_*. = 15, pooled 3 experiments) and 48 hours (n_PBS_ = 6, n*_S.pn._* = 10, pooled 2 experiments) post-infection (hpi), and following (**C**) high tidal volume (HVT, n = 11) mechanical ventilation or in non-ventilated control mice (NV ctr, n = 8). (**D**, **E**) Murine CXCL-chemokine concentrations measured in cell culture supernatants 24 h post infection or stretch by ELISA in (**D**) T7 – murine AECII cell line following *S.pn.* infection (MOI 0.1, n = 3) or 18% stretch (n = 2), and (**E**) isolated primary murine lung EpiCs and AMs following *S.pn.* infection (MOI 1, n = 3), representative of 2 – 3 experiments. (**F**) Immunohistochemistry of *S.pn.*- or PBS-infected murine lungs displaying nuclear β-galactosidase fluorescence (AF488, arrows) in BEC and AEC of *Cxcl5*^−/−^ mice (transgenic for *LacZ*) and WT controls. Scale bar indicates 50 µm. Images representative of n = 5 animals from 2 experiments. (**G, H**) Human CXCL5 chemokine concentrations measured in cell culture supernatants 24 hours post-infection or stretch by ELISA in (**G**) BEAS-2B BEC line following *S.pn.* infection (MOI 0.1, n = 9, pooled from 3 experiments) and (**H**) HPAECs following *S.pn.* infection (MOI 1, n = 8, pooled from 4 experiments) or 18% stretch (n = 9, pooled from 4 experiments). (**A – F, G, H**) Mean ± SEM. (**A, B, D – F**) Two-way ANOVA, Šídák’s multiple comparisons test, (**C, G, H**) Mann Whitney test, unpaired. * *P* < 0.05, ** *P* < 0.01, *** *P* < 0.001 and **** *P* < 0.0001. BALF, Bronchoalveolar lavage fluid; CAP, community acquired pneumonia; AEC, alveolar epithelial cells; II, type 2; BEC, bronchial epithelial cells; EpiC, epithelial cells; AM, alveolar macrophages; MOI, multiplicity of infection; h, hours.

In a murine ATII cell line (T7), significantly enhanced CXCL1 and CXCL5 concentrations were detected following *S.pn.* infection or mechanical stretch (Figure 1D). Isolated murine-derived primary lung epithelial cells demonstrated enhanced CXCL-chemokine secretion upon *S.pn.* infection, whilst primary murine alveolar macrophages (AM) produced CXCL1 and CXCL2, but not CXCL5 (Figure 1E). Production of CXCL5 by murine primary bronchial epithelial cells (BECs) and alveolar epithelial cells (AECs) upon *S.pn.* infection was also confirmed by immunohistochemistry, as depicted by the nuclear localization signal of β-galactosidase (LacZ), which was used to replace the *Cxcl5* gene in *Cxcl5*^−/−^ mice (Figure 1F) [13]. Furthermore, hCXCL5, hCXCL6 and hCXCL8 production was induced by pneumococcal infection in a human BEC cell line (BEAS-2B) (Figure 1G, E1A). Human primary alveolar epithelial cells (HPAECs, Cell Biologics, Chicago, IL, USA) secreted hCXCL5 following *S.pn.* infection as well as cyclic stretch stimulation (Figure 1H, E1B).

### CXCL5 deficiency results in increased pneumococcal burden without affecting survival

Next, we probed how absence of CXCL5 affects disease manifestation. During the course of pneumococcal pneumonia, WT and *Cxcl5*^−/−^ mice exhibited no significant differences in survival or changes in body temperature. Weight loss was transiently greater in WT mice at 60 and 72 hpi, at a time were mice reached humane end point criteria, but not 24 hpi (Figure 2A–B, E1C–D).

**Figure 2.**
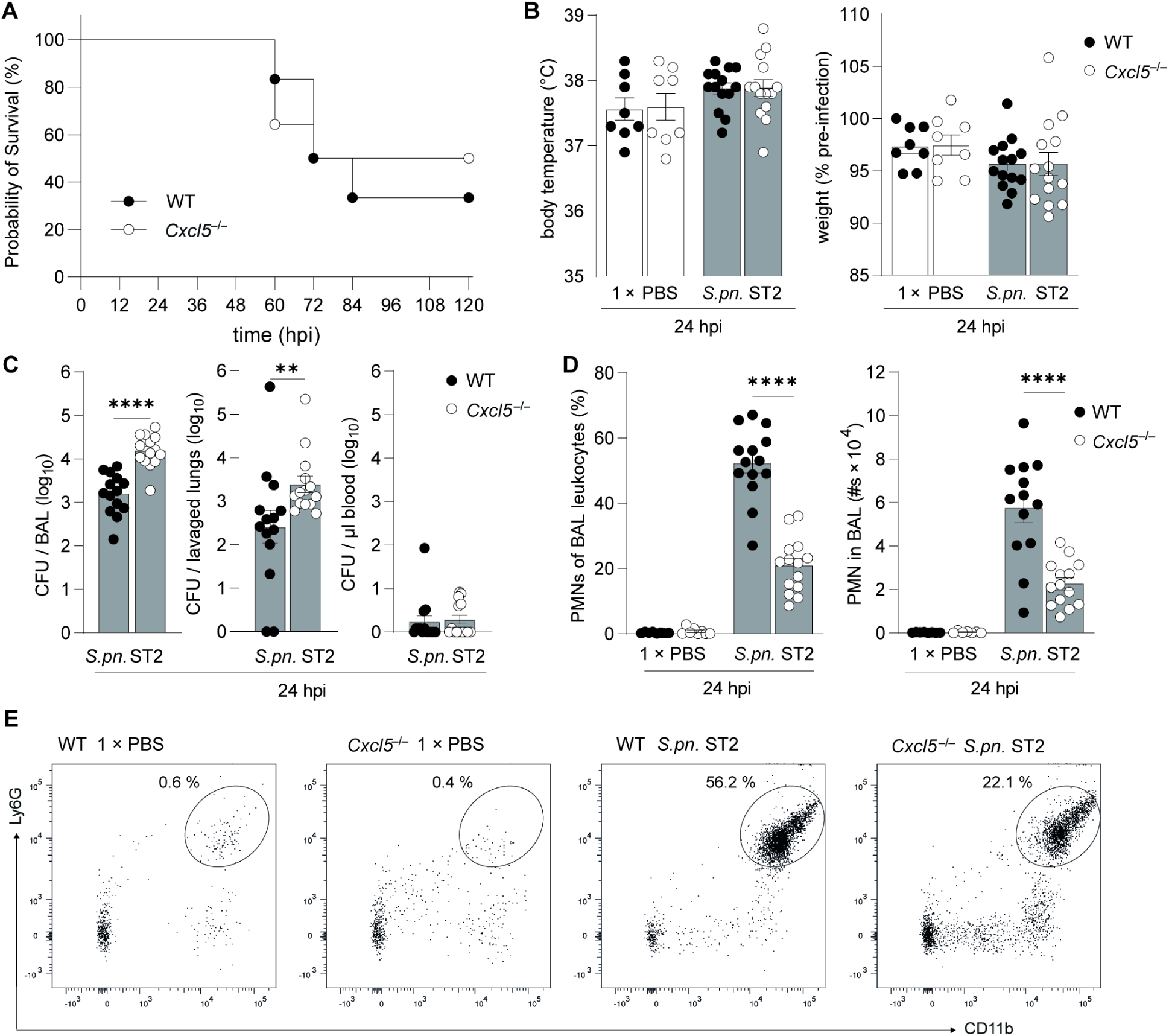
Absence of CXCL5 impairs pulmonary bacterial clearance without altering survival of mice following infection. WT and *Cxcl5*^−/−^ mice were intranasally infected with 5 x 10^6^ CFU *S.pn.* ST2. (**A**) Survival (%) of WT (n = 12) and *Cxcl5*^−/−^ (n = 14) mice up to 120 hpi, data pooled from 2 experiments. Mice indicated as non-surviving had been euthanized due to reaching of humane end point criteria before developing severe symptoms. Kaplan-Meier curves, log-rank (Mantel-Cox) test. (**B** – **E**), 24 hpi time point analysis of *S.pn.* ST2 infected WT (n = 14), *Cxcl5*^−/−^ (n = 14) and PBS ctr WT (n = 8) and *Cxcl5*^−/−^ (n = 8) mice, data pooled from 2–3 experiments, mean ± SEM. (**B**) Infection-induced changes in body temperature and weight of WT and *Cxcl5*^−/−^ mice. (**C**) Bacterial burden quantified in BAL (per ml), lavaged lungs and blood (per µl) of WT and *Cxcl5*^−/−^ mice. Mean ± SEM. Mann-Whitney test. (**D**) Frequencies and total numbers of BAL neutrophils in *S.pn.*- or PBS-infected WT and *Cxcl5*^−/−^ mice. Mean ± SEM. One-way ANOVA, Šídák’s multiple comparisons test, WT vs *Cxcl5*^−/−^ mice. (**D**, PMN #s: one outlier identified by ROUT method and removed WT = 2.17 x 10^5^ cells). (**E**) Representative dots blots of BAL PMNs. *** *P* < 0.001 and **** *P* < 0.0001. BAL, Bronchoalveolar lavage; hpi, hours post infection; WT, wild type; CFU, colony forming units; *S.pn.*, *Streptococcus pneumoniae;* PMN, neutrophils.

At 24 hpi, significantly higher bacterial loads in the BAL and lungs of *S.pn.*-infected *Cxcl5*^−/−^ mice were recorded. This coincided with reduced numbers of alveolar neutrophils as well as altered numbers of other leukocytes (Figure 2C–E, E2B).

### Absence of CXCL5 improves alveolar barrier permeability independent of neutrophil recruitment

Histopathological evaluation of *S.pn.*-infected murine lungs revealed a similar degree of pathology between WT and *Cxcl5*^−/−^ mice (Figure 3A). The overall inflammation score (including purulence, catarrhal purulence, perivascular edema, pleuritis and steatitis), remained balanced between *S.pn.*-infected lungs of WT and *Cxcl5*^−/−^ mice (Figure 3B). Furthermore, both WT and *Cxcl5*^−/−^ mice displayed predominantly interstitial, partly histiocytic, partly purulent pneumonia (Figure 3A). When assessing inflammatory cytokine and chemokine secretions into BALF post-*S.pn.* infection, a generally similar profile of protein concentrations could be detected across WT and *Cxcl5*^−/−^ mice, except for VEGF and CXCL5; which were higher in WT mice (Figure 3C). In plasma, only CXCL5 concentrations were different (Figure 3D). Notably, despite higher bacterial burden, reduced neutrophil recruitment and no differences in inflammatory mediators on a protein level, lung permeability was significantly lower in *S.pn.*-infected *Cxcl5*^−/−^ mice compared to WT mice 24hpi (Figure 3E). Similarly, in a murine model of VILI, absence of CXCL5 resulted in reduced neutrophil recruitment and preservation of alveolar-capillary barrier integrity after 4 hours of ventilation (Figure E3A–C). In order to better understand the impact of CXCL5 on lung barrier permeability independent of neutrophil chemotaxis, we turned to an isolated-perfused mouse lung (IPML) model. CXCL5 binds heparan sulfate with high affinity [20], and accordingly, we found CXCL5 localized in proximity to the epithelial glycocalyx under steady state (Figure 3F). Having recently demonstrated that the glycocalyx can be disrupted by exposure to pneumolysin [21], we anticipated local CXCL5 release by pneumolysin-induced glycocalyx disruption. Shortly (30 min) following intravenous (i.v.) or intratracheal (i.t.) pneumolysin instillation of WT and *Cxcl5*^−/−^ mice, we measured the impact on the lung barrier by assessing the amount of transvasated HSA into the BALF in an IPML model. Application of pneumolysin from the vascular side moderately opened the lung barrier, with no impact of *Cxcl5* deletion. However, when applied intratracheally, the lung barrier remained more intact in *Cxcl5*^−/−^ mice independent of neutrophil recruitment (Figure 3G).

**Figure 3.**
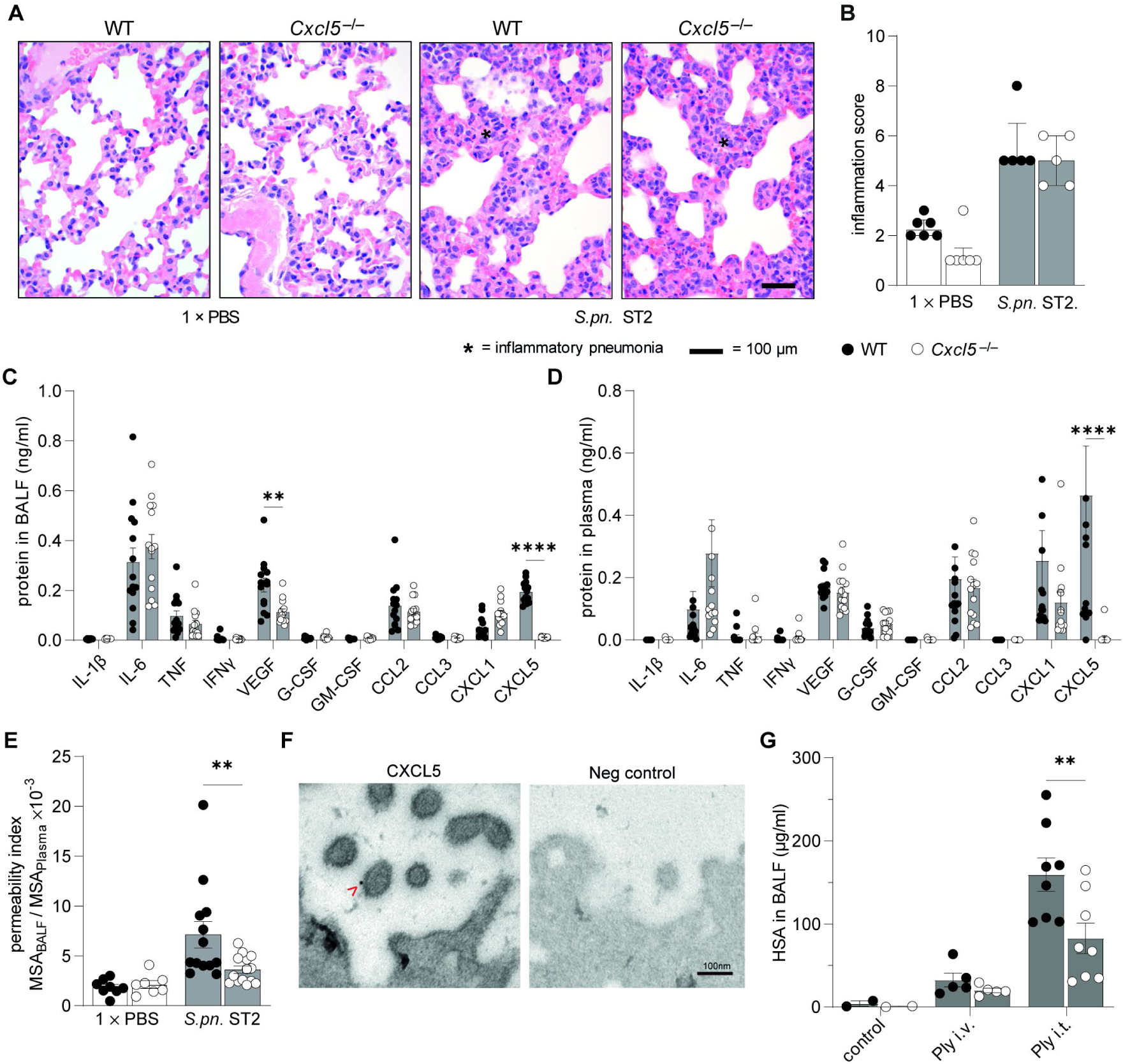
Lack of CXCL5 improves lung barrier function in mice. (**A**, **B**) Histopathological 24 hpi time point analysis of *S.pn.* ST2 infected WT (n = 5), *Cxcl5*^−/−^ (n = 5) and PBS ctr WT (n = 6) and *Cxcl5*^−/−^ (n = 6) mice. (**A**) representative H & E staining (scale bar 100 µm) and (**B**) inflammation score data pooled from 2 experiments. (**C** – **E**) 24 hpi time point analysis of *S.pn.* ST2 infected WT (n = 13 – 14), *Cxcl5*^−/−^ (n = 14) and (**E**) PBS ctr WT (n = 8) and *Cxcl5*^−/−^ (n = 7) mice, data pooled from 2 – 3 experiments. Protein levels of inflammatory mediators measured in (**C**) BALF and (**D**) plasma of infected WT and *Cxcl5*^−/−^ mice. (**E**) Ratios of mouse serum albumin (MSA) in BALF to plasma of *S.pn.* -infected mice reflecting lung barrier integrity, calculated from ELISA readouts. (**F**) Transmission electron microscopy (TEM) of naïve WT murine lung tissue displaying CXCL5 staining on surface of alveolar epithelial cells. Arrow indicates 10 nm gold-particle CXCL5-staining on alveolar microvilli surface. Representative images (n = 3). Scale bar 100 nm. (**G**) Concentration of HSA in BALF of IPML of WT (ctr n = 2; Ply i.v. n = 5; Ply i.t. n = 8) and *Cxcl5*^−/−^ (ctr n = 2; Ply i.v. n = 5; Ply i.t. n = 8) mice. (**A**) asterisk indicates inflammatory pneumonia. (**B** – **E**; **G**) Mean ± SEM. One-way ANOVA, Šídák’s multiple comparisons test, WT vs *Cxcl5*^−/−^ mice. ** *P* < 0.01 and **** *P* < 0.0001. *S.pn.*, *Streptococcus pneumoniae*; WT, wild type; hpi, hours post infection; MSA, mouse serum albumin; TEM, transmission electron microscopy; BALF, Bronchoalveolar lavage fluid; HSA, human serum albumin; IPML, isolated perfused mouse lungs; Ply, Pneumolysin; i.v., intravenous; i.t., intratracheal.

### Transcriptomic profiling of alveolar epithelial and endothelial cells in the absence of CXCL5

We performed spatial transcriptomics on lung tissue from a *S.pn.* ST3-infected *Cxcl5*^−/−^ compared to an infected WT mouse and a PBS control (ctr) WT mouse (Figure 4A, E4A). Analysis of differentially expressed genes (DEG) between infected WT and *Cxcl5*^−/−^ mice revealed upregulation of inflammatory genes (e.g. *Cxcl1*, *Cxcl10*, *Il6*) in the infected WT mouse, with highest expression in peribronchial areas. *Cxcl5* expression was observed specifically in peribronchial areas close to larger airways, limited to the infected WT mouse (Figure 4B–D, E4B–C).

**Figure 4.**
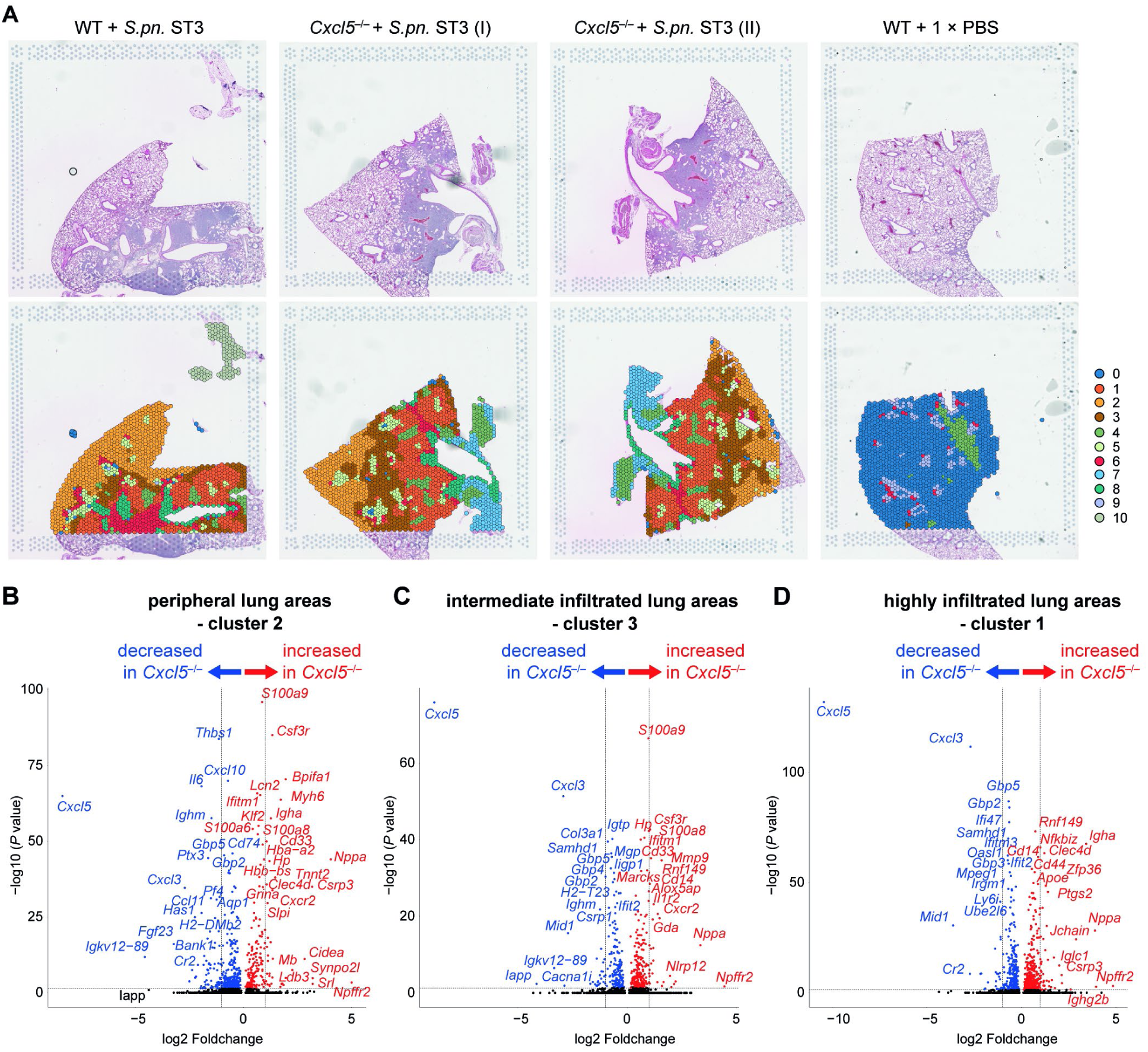
Spatial sequencing allows for topological analysis of infection induced gene expression in WT and *Cxcl5*^−/−^ mice. (**A**) H&E-stained lung sections from *S.pn.* ST3 infected WT (n = 1), *S.pn.* ST3 infected *Cxcl5*^−/−^ (n = 1, two sections) and PBS ctr WT (n = 1) mice used for spatial transcriptomics analysis (upper row) and unbiased clustering based on gene expression (lower row). Unsupervised clustering identified 11 distinct clusters which matched with the following histopathological areas: intact uninfected peripheral areas (cluster 0), intact infected peripheral areas (cluster 2), intermediate (cluster 3) and highly infiltrated areas (cluster 1) as well as edematous areas (cluster 6). Volcano plots displaying differentially expressed genes of histologically intact (**B**) peripheral lung areas (cluster 2), (**C**) intermediate infiltrated lung areas (cluster 3) and (**D**) highly infiltrated lung areas (cluster 1) of *S.pn*. ST3-infected WT mice compared to infected *Cxcl5^−/−^* mice. Differential gene expression analysis was performed using a Wilcoxon rank-sum test, displayed *P* values were adjusted for multiple testing using the Bonferroni correction. Dashed lines indicate adjusted *P* value of 0.05 (horizontal) or log2-fold changes of −1 and 1 (vertical).

Next, we performed single-cell transcriptomics focusing on barrier-forming cells and subclustered AT2 and EC, respectively, in *S.pn.* ST3-infected or PBS ctr WT and *Cxcl5*^−/−^ mice (n=3, all groups) (Figure E5A–C). Although AT2 displayed limited DEG between *S.pn.* ST3-infected *Cxcl5*^−/−^ and WT mice, a distinct pattern of genes in *Cxcl5*^−/−^ mice were upregulated, which have been described to be involved in establishment and maintenance of epithelial barriers (*Vcl*, *Stxbp6*, *Egfr*, *Lrg1*) (Figure 5A, E6A–B; full DEG list in supplementary data). Collectively, these upregulated genes may influence cellular processes associated with cell adhesion and migration, vesicle trafficking and cell proliferation and are partly linked to actin cytoskeletal remodeling [22–25]. Gene set enrichment analysis confirmed corresponding activated pathways in *Cxcl5*-deficient mice, including “cell junction organization”, “cell migration” and “cell motility” within the top 10 activated pathways in AT2 cells (Figure 5B). DEG on EC indicated upregulation of inflammatory genes primarily contributing to the JAK/STAT pathway such as *Stat2*, *Irf1*, *Irf7* in infected WT mice (Figure 5C, E6C–D). In line with the increased VEGF levels in BALF of infected WT mice (Figure 3C), we detected increased expression of *Vegfa* in EC of WT mice. Additionally, *Tjp1* was downregulated in EC, which might contribute towards endothelial barrier alterations in infected WT animals. However, other observed trends in barrier-related genes did not reach statistical significance in epithelial or EC (Figure E6B, D). Gene set enrichment analysis in EC supported these observations highlighting enhanced expression of “innate immune response”, “defense response” and “response to cytokine” in EC from WT compared to *Cxcl5*^−/−^ mice (Figure 5D). Cell-cell communication analysis highlighted the role of crosstalk between structural cells of the alveoli and neutrophils in lungs during infection. In *Cxcl5*^−/−^ mice, AT2 cells showed increased ligand-receptor interactions with themselves, fibroblasts, macrophages and dendritic cells (DCs) as well as EC. Conversely, WT mice showed increased EC interactions with fibroblasts, neutrophils, T- and NK cells as well as macrophages and DCs (Figure E7A). Expressed ligands released by neutrophils were predicted to induce DEG predominantly in WT EC, whilst ligands from structural cells (AT2, endothelial, fibroblasts) induced a broader pattern of DEG in WT and *Cxcl5*^−/−^ neutrophils (Figure E7B). In *Cxcl5*^−/−^ mice, this included upregulation of several genes encoding metallothioneins, which have been reported to reduce oxidative stress and neutrophilic lung inflammation in LPS-induced ALI in addition to protection of vascular integrity [26]. In WT mice, interferon response genes *IFIT2*, *IFIT3*, and *IFI27* were upregulated, in line with earlier results indicating heightened inflammation levels in WT EC.

**Figure 5.**
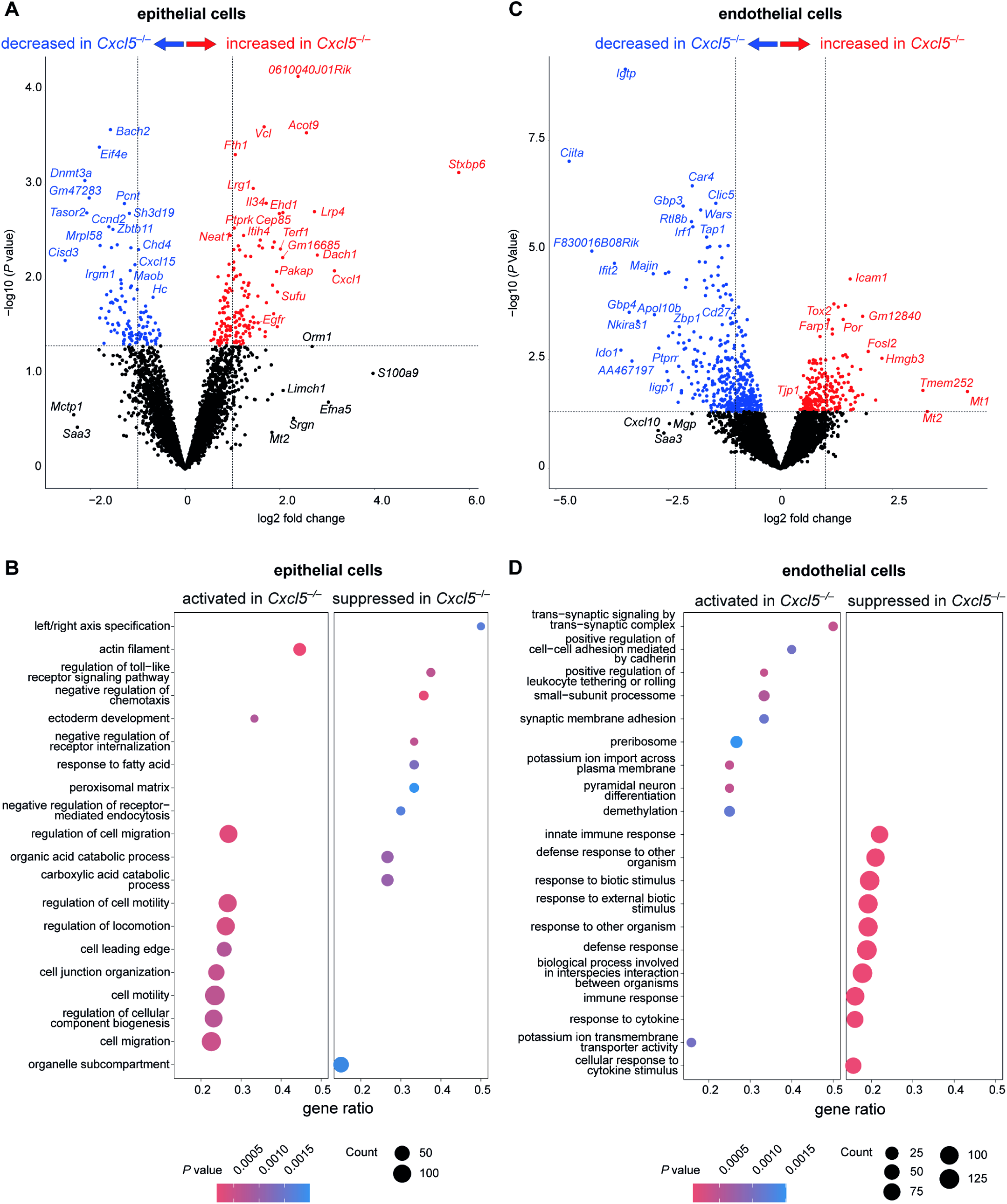
Absence of CXCL5 results in higher expression of epithelial barrier genes upon infection. (**A**) Volcano plot displaying differentially expressed genes of AT2 cells between *S.pn*. ST3-infected WT and *Cxcl5*^−/−^ mice. (**B**) Gene Set Enrichment Analysis of AT2 cells (*S.pn.* ST3-infected) based on GO sub-ontologies Biological Process (BP), Cellular Component (CC) and Molecular Function (MF), were performed using clusterProfiler. (**C**) Volcano plot displaying differentially expressed genes of endothelial cells between *S.pn*. ST3-infected WT and *Cxcl5*^−/−^ mice. (**A**, **C**) Colored dots indicate differential expressed genes with *P* values below 0.05. Dashed lines indicate *P* value of 0.05 (horizontal) or log2-fold changes of −1 and 1 (vertical). (**D**) Gene Set Enrichment Analysis of endothelial cells (*S.pn.* ST3-infected) based on GO sub-ontologies Biological Process (BP), Cellular Component (CC) and Molecular Function (MF). Differential gene expression analyses were performed using the edgeR framework.

### Human CXCL5 induces epithelial but not endothelial barrier failure in response to TNF priming

To better understand the differential effects of *Cxcl5*-deficiency on gene expression and barrier integrity, we conducted an impedance-based *in vitro* assay (ECIS^®^) to quantify changes in monolayer resistance following CXCL5 exposure. HPAECs and human pulmonary microvascular endothelial cells (HPMEC) were primed with TNF and then exposed to hCXCL5 (5-78aa or 8-78aa isoforms).

ECIS^®^-based barrier quantification of HPMEC monolayers revealed that neither form of hCXCL5 had an additional effect on TNF-induced endothelial barrier disruption (Figure 6A, E8A). In contrast, treatment of TNF-primed HPAECs with the short-length hCXCL5(8-78aa) enhanced disruption of epithelial barrier integrity, whereas this additive effect was absent with the less potent long-length hCXCL5(5-78aa) (Figure 6B, E8B). In line with these findings, immunofluorescence staining of HPMECs showed gap formations after 3h of TNF-priming, independent of hCXCL5 treatment, with gaps persisting up to 12h, indicating increased endothelial cell death and reduced recovery (Figure 6C). In contrast, TNF-priming alone had no effect on HPAECs, but when combined with short-length hCXCL5, transient gap formation was observed at 3h, which resolved by 12h (Figure 6D).

**Figure 6:**
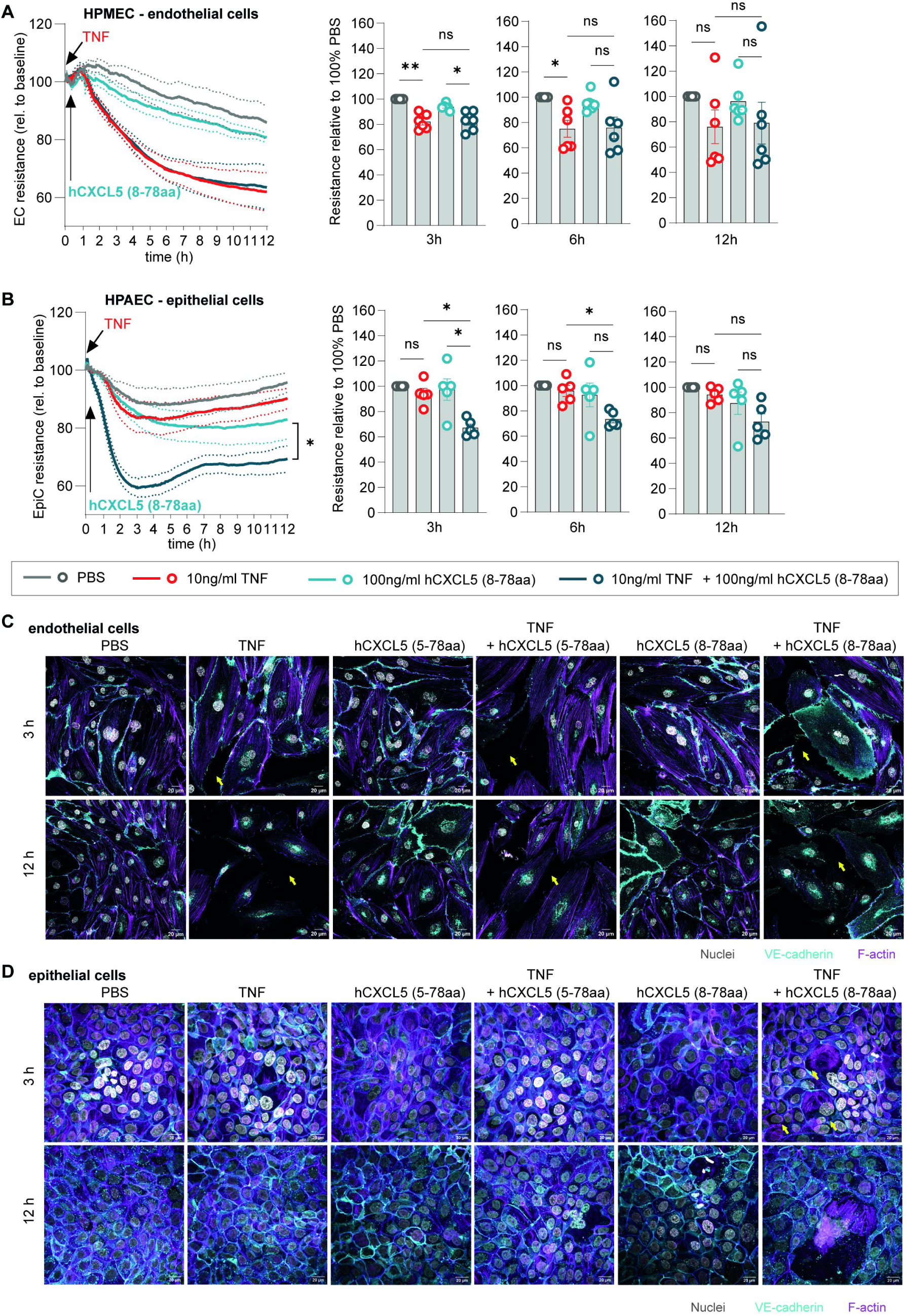
Human CXCL5(8-78aa) induces barrier failure in TNF-primed human primary alveolar epithelial cells. Human primary cells were primed with 10 ng/ml TNF followed by stimulation with 100 ng/ml hCXCL5 (5-78aa or 8-78aa) 10 min later. Line plots show the quantified barrier integrity of HPMEC (**A**) and HPAECs (**B**) by electric cell impedance sensing (ECIS^®^) over 12h and scatter dot plots indicate the percent changes in electrical resistance of primed and stimulated cells relative to control (PBS)-treated cells 3h, 6h and 12h following stimulation. HPMEC (**C**) and HPAEC (**D**) monolayers primed and treated for 3h and 12h under the same conditions were stained with VE-Cadherin-(FITC), E-Cadherin (cyan), phalloidin-AF546 (magenta) and DAPI (grey), followed by visualization by immunofluorescence. (**A-B**) Data are presented as mean ± SEM; for curve analysis, two-way ANOVA, Dunnett’s multiple comparisons test, * *P* < 0.05 or for comparison of normalized resistance values at the designated timepoints, Kruskal Wallis-test was performed; *P* values indicated in Figure. (**C-D**) Yellow arrows indicate gap formation. Scale bars, 20µm.

## DISCUSSION

In this study we show that CXCL5 increases barrier permeability during pneumonia and VILI. Elevated CXCL5 concentrations were detected in BALF of patients with severe CAP and HPAECs following *S.pn.* infection or cyclic stretch. In murine *S.pn.* pneumonia and VILI, absence of CXCL5 reduced alveolar neutrophil infiltration and improved lung barrier function, by direct effects of CXCL5 on the epithelial barrier independent of neutrophil recruitment. This finding was further substantiated by experimental *ex vivo* and *in vitro* studies of the lung barrier.

CXCL-chemokines are essential for neutrophil chemotaxis to infected tissue, however the contributions of individual CXCL ligands towards pneumonia outcome are distinct. For instance, genetic deletion of *Cxcl1* culminates in increased mortality of mice following *S.pn.* infection due to insufficient neutrophil recruitment and inability to contain bacterial growth [27], which is in contrast to our findings in *Cxcl5*-deficient mice. The complex role of neutrophils in pneumococcal pneumonia was further illustrated in a study where neutrophils were depleted before or 18h after *S.pn.* infection, resulting in sharply elevated bacterial loads and increased mortality in pre-depleted mice compared to improved survival of mice with neutrophils depleted 18hpi [28]. Correspondingly, neutropenic individuals are a major risk group for fatal bacteremic pneumonia [29]. This notion is further supported by a model of cecal ligation and puncture (CLP)-induced sepsis, in which negative regulation of CXCL5 by growth and differentiation factor-15 (GDF15) results in impaired neutrophil recruitment, leading to enhanced bacterial burden and increased mortality [30]. Nevertheless, transepithelial migration of neutrophils may disrupt epithelial barrier function [31] and extended neutrophilic inflammation coincides with increased lung permeability during ALI [32]. A recent study elucidated that anti-Ly6G-mediated neutrophil reduction combined with antibiotics significantly reduced lung barrier permeability and edema compared to antibiotics alone, despite higher bacterial burden in BAL [33]; which is concurrent with our findings. Based on our findings, targeting CXCL5 alongside antibiotics could enhance treatment by improving lung barrier function and reducing neutrophilic inflammation, surpassing the effectiveness of antibiotics alone.

Established hallmarks of ALI include increased vascular permeability and pulmonary edema [4, 5], making these pathological processes major candidates for therapeutic strategy development. Targeting the endothelial barrier has shown therapeutic promise during severe pneumonia [34], however clinical studies investigating the lung epithelial barrier remain lacking. Recent studies demonstrate contribution of glycosaminoglycans to lung epithelial barrier function during pneumococcal pneumonia and ALI, albeit lacking specific molecular mechanisms [5, 35]. Hence, improved molecular understanding of how the lung epithelial barrier is regulated remains imperative for novel therapy development. The CXCL-CXCR2 axis’ potential as a targeting point for clinical studies on the treatment of CAP and ALI have been highlighted in literature and examined in various clinical trials. Despite having been well tolerated in a phase II randomized controlled study in hospitalized patients with COVID-19 (NCT04794803) [12], antagonizing CXCR2 has also proven to be problematic in other clinical trials. In a randomized, double-blinded, placebo-controlled phase II study in patients with moderate to severe COPD (NCT01006616), treatment with a CXCR2 antagonist resulted in significant improvement of FEV1 and numerical improvement in COPD score in current smokers compared to placebo treatment, however, dose-related study discontinuations were recorded due to absolute neutrophil count decreases [36]. In another randomized, double-blinded, placebo-controlled phase IIa proof-of-concept study in patients with bronchiectasis (NCT01255592), therapy with a CXCR2 antagonist reduced sputum neutrophil counts, but did not improve clinical outcomes and even led to treatment discontinuations due to adverse events including lower respiratory tract infections, pneumonia and infective exacerbation of bronchiectasis [37]; underscoring the limitations of antagonizing CXCR2 in the context of certain lung pathologies. Owing to the multiple CXCL-chemokine ligands of CXCR2 responsible for neutrophil chemotaxis [38], targeting individual CXCL ligands may offer an advantage over CXCR2-targeted therapeutics: Neutrophil recruitment is not entirely abolished, but beneficially tempered, as demonstrated in our murine acute lung injury model using *Cxcl5*-deficient mice. In fact, an *in vitro* study revealed that during exposure of neutrophils to plasma of patients with COVID-19, the IL-8-CXCR1/2-axis can be effectively targeted not just by CXCR1/2 antagonization, but also through IL-8 inhibition in order to reduce neutrophil activation [39]. Hence, considering that not just IL-8, but also CXCL5 levels were consistently correlated to neutrophil concentrations in lung fluids of patients with ARDS [40], targeting CXCL5 in a clinical setting could offer therapeutic value. Taken together, we demonstrate a novel role for CXCL5 in inducing lung epithelial barrier permeability, warranting further clinical validation of CXCL5 chemokine antagonization as next-generation therapeutic in pneumonia.

## ACKNOWLEDGEMENTS

The authors would like to thank Stephanie Hübl, Charlene Lamprecht and Cornelia Zieger for assistance in histopathology and Farzin Mashreghi and Gabriela Guerra as well as Jeannine Wilde, Kirsten Richter and Madlen Sohn for RNA sequencing. Pneumolysin was a kind gift from Timothy J. Mitchell. The funders had no role in study design, data collection, data analysis, interpretation, or writing of the manuscript.

## Author Contributions

Conceptualization: G.N.; methodology: S.B., C.G., P.P., B.G., L.M., K.H., E.L.-R., V.G., S.-M.W., U.B., A.T., K.D., H.K., S.K., K.M., T.H., S.M.V., S.W., G.N.; software/data analysis: S.B., C.G., P.P., B.G., L.M., K.H., E.L.-R., V.G., S.-M.W., A.T., K.D., H.K., K.M., G.N.; Resources: B.G., T.H., S.V., A.D.G., W.M.K., L.E.S., CAPSyS Study group; writing—original draft preparation: S.B., C.G., P.P. and G.N.; writing—review and editing: all; supervision: S.-M.W., B.G., A.D.G., W.M.K., L.E.S., M.W., G.N.; project administration: S.B., C.G., G.N.; funding acquisition: M.W. and G.N.; All authors have read and agreed to the published version of the manuscript.

This research was supported by the German Federal Ministry of Education and Research, Germany (BMBF) grants e:Med CAPSyS - 01ZX1604B, e:Med SYMPATH - 01ZX1906A and MAPVAP - 01KI2124, by the German Center for Lung Research (Deutsches Zentrum für Lungenforschung, DZL) grants PROGRESS - 82DZLJ19B1 and funded by the Deutsche Forschungsgemeinschaft (DFG, German Research Foundation) – Project ID 431232613 – SFB 1449 and project ID 114933180 – SFB-TR84.

