## Supplement for "Neutrophil-chemoattractant CXCL5 induces lung barrier permeability in acute lung injury"

#### EXTENDED MATERIAL AND METHODS

##### Chemokine analysis of human BAL

Our data includes a control group of 5 patients without apparent lung disease (non-CAP patients) and 6 patients with community-acquired pneumonia (CAP patients) who committed to the CAPSyS-Project recruitment protocol and had confirmed mono-dominated *S.pn.* infection by 16S rRNA amplicon sequencing and not received antibiotic treatment prior hospitalization (personal communication T. Hain on behalf of CAPSyS study group).
Chemokine protein profiling was performed by diluting patient BAL fluid (BALF) samples 1:10 in 10x protease inhibitor (Roche, Basel, Switzerland) to obtain 1x concentration in BALF (45 µl BALF fluid + 5 µl 10x protease inhibitor). The chemokines CXCL8 (IL-8), hCXCL5 (ENA-78) and hCXCL6 (GRO-α) were analyzed using bead-based immunoassays – custom-modified LEGENDplex™ human anti-virus response and pro-inflammatory chemokine panels (BioLegend, San Diego, CA, USA) – according to the manufacturer's instructions. Samples were acquired on a BD Canto II equipped with FACSDiva software (BD Bioscience, Heidelberg, Germany) and analyzed with LEGENDplex™ data analysis software [1]. Values below the limit of detection (LOD) lowest standard (C1) were shown as C1/2.

##### Dissection and sampling of experimental animals

##### Murine bronchoalveolar lavage (BAL)

To analyse BAL bacterial burden, protein and cells, airways were washed twice with 0.8 ml 1 × PBS supplemented with a protease inhibitor cocktail (Roche, Basel, Switzerland). BAL serial dilutions were plated for enumeration of bacterial burden onto blood agar plates. The BAL suspension was centrifuged at 470 × g for 5 min, and the supernatant (BALF) frozen at -80 °C until analysis. Cell pellets were resuspended in 1 × PBS, containing 0.2 % fetal calf serum (FCS) and subjected to flow cytometry analysis.

##### Murine blood samples

Blood samples for bacterial burden and plasma protein analysis were taken from the vena cava (or arteria carotis in ventilation experiments) and collected in EDTA-coated tubes. Blood serial dilutions were plated for enumeration of bacterial burden onto blood agar plates. For plasma analysis, EDTA-blood was centrifuged at 2.000 × g for 10 min, plasma collected and stored at -80 °C until analysis.

**Murine lung homogenates**

To assess lung bacterial burden, lungs were perfused with 1 × PBS. Lung lobes were dissected, immediately dispersed in 1 × PBS supplemented with a protease inhibitor cocktail and homogenized in gentleMACS separator M tubes (Miltenyi Biotec, Bergisch Gladbach, Germany). Serial dilutions were plated for enumeration of bacterial burden onto blood agar plates (BD Bioscience, Heidelberg, Germany).

**Histopathology and Immunohistochemistry of murine lungs**

Lungs were immersion-fixed for 24 h in 10 % formalin and routinely embedded in paraffin. Sections were cut at 2 µm thickness, dewaxed and stained with haematoxylin–eosin. Samples were analyzed by veterinarian pathologists independently and blinded to the study groups. The extent of lung inflammation and damage was quantified by a severity score ranging from 0 to 5 (0, no inflammatory cells; 1, scattered inflammatory cells; 2, few inflammatory cells; 3, moderate numbers of inflammatory cells with multifocal scant exudate; 4, large numbers of inflammatory cells with moderate amounts of exudate and multifocal necrosis; and 5, widespread necrosis and large numbers of inflammatory cells with abundant exudates).

In *Cxcl5*$^{-/-}$ mice the *Cxcl5* gene was replaced with a *β-galactosidase* (*LacZ*) gene containing a nuclear localization signal sequence [2]. Nuclear β-galactosidase was revealed by immunohistochemistry after heat-mediated antigen retrieval using a purified chicken antibody polyclonal to ß-galactosidase (1:5000, ab9391, abcam, Cambridge, UK) at 4 °C overnight. Subsequently, slides were incubated with biotinylated, secondary goat anti-chicken IgG antibody (1:200, BA-9010, Vector, Burlingame, CA, USA) and HRP-coupled streptavidin. Diaminobenzidine (DAB) was used for color development. Slides were counterstained with hematoxylin and coverslipped. For immunofluorescence analysis slides were incubated with the ß-galactosidase antibody (1:200) overnight at 4 °C as described above and with Alexa Fluor 488-conjugated secondary goat anti-chicken IgY antibody (1:200, ab150169, abcam, Cambridge, UK) for 45 min at room temperature and mounted with Roti-Mount FluorCare DAPI (4,6-diaminidino-2-phenylindole) (Carl Roth, Karlsruhe, Germany). Adequate negative controls were conducted. All slides were analyzed and pictures were taken with an Olympus BX41 microscope equipped with a DP80 camera (Olympus, Tokyo, Japan).

**Electron microscopy of lungs**

Lungs of WT mice that were isolated and perfused with 1.5% formaldehyde solution + 1.5% glutaraldehyde (GA) in 0.15 M HEPES were cut into small pieces and the fixative solution was changed to 4% formaldehyde solution + 0.1% GA in 0.2 M HEPES for post-embedding immuno-labeling. After overnight fixation at 4°C, the pieces were infiltrated in 2.3 M sucrose in

PBS, frozen in liquid nitrogen and transferred to an automated freeze substitution system (AFS2, Leica, Wetzlar, Germany). Freeze substitution was performed at -90°C using methanol containing 0.5% uranyl acetate. Subsequently, samples were embedded in Lowicryl HM20 at -45°C [3] and ultrathin sections of 70 nm were cut. The nickel grids were floated with ultrathin sections side facing a drop of the following solution: 50 mM glycine (Carl Roth, Karlsruhe, Germany), 0.1% BSA-c (Aurion, Wageningen, Netherlands) in PBS (pH 7.4) for 20 min and subsequently blocked for 30 min using blocking solution (Aurion, Wageningen, Netherlands). Immuno-labeling was performed by floating the grid on 25 µl drops of specific anti-CXCL5 antibody (1:100, PA5-115069, Invitrogen, Waltham, MA, USA) on parafilm in a wet chamber at 4°C overnight. After washing in incubation buffer (0.1% BSA-c in PBS), bound antibody was detected following incubation with a 10 nm-gold-conjugated goat-anti-rabbit antibody (Aurion, Wageningen, Netherlands), at a final dilution of 1:30 in incubation buffer for 90 min at room temperature. Finally, the sections were rinsed in incubation buffer (six times) and PBS (three times), postfixed with 2.5% GA in PBS and contrasted with an aqueous solution of 1% phosphotungstic acid (15 min) followed by 4% uranyl acetate (15 min). Lastly, grids were washed three times in distilled water. Negative controls were performed without primary antibody but incubation with the 10 nm-gold-conjugated goat-anti-rabbit antibody (Aurion, Wageningen, Netherlands) to rule out unspecific binding of the secondary antibody.
Visualization was performed by using a Zeiss Leo 906 electron microscope at 80 kV acceleration voltage and equipped with a slow scan 2K CCD camera (TRS, Moorenweis, Germany).

**Flow cytometry**

For flow cytometry of innate immune cells, isolated cells were stained with anti (α)-Ly6G (1A8, BD Biosciences, Heidelberg, Germany), αCD11b (M1/70, BD Biosciences, Heidelberg, Germany), αF4/80 (BM8, eBioscience, San Diego, CA, USA), αLy6C (AL-21, BD Biosciences, Heidelberg, Germany), αMHCII (M5/114.15.2, Thermo Fisher Scientific, Waltham, MA, USA) and αCD11c (HL3, BD Biosciences, Heidelberg, Germany) antibodies. The FL-1 channel was used to determine autofluorescence of cells (gating strategy in Figure E2). All stained cells were acquired using a BD FACS Canto II and analysed with BD FACSDiva (BD Biosciences, Heidelberg, Germany) and FACS-Analyzer software. Total cell numbers were calculated using CountBright™ Absolute Counting Beads (Thermo Fisher Scientific, Waltham, MA, USA) according to the manufacturer’s instructions.

**Multiplex and ELISA on murine cytokines/chemokines**

Samples from BALF, plasma and cell culture supernatants were thawed and appropriately diluted. Measurements were performed using multiplex bead-based immunoassay kits (ProcartaPlex®, Thermo Fisher Scientific, Waltham, MA, USA) or ELISA (R&D, Minneapolis, MN, USA) according to manufacturers' instructions. Samples were acquired on a Bio-Rad instrument (Hercules, CA, USA) or SpectraMax ELISA Reader (Molecular Devices, San José, CA, USA).

**Single-cell barcoding, cDNA, library preparation and sequencing**

Single-cells from lung tissue were isolated as described previously [4]. Antibody-based hashtag oligos for sample multiplexing were used according to manufacturers' instructions (TotalSeq™-B, BioLegend, San Diego, CA, USA). Labeled cells were filtered through a 40 µm Flowmi® cell strainer (Sigma-Aldrich, St. Louis, MO, USA) and counted. Pooled cells from six samples each were adjusted to a final concentration of ~1,200 cells/ml or ~1,500 cells/ml and split into two equal pools prior to loading onto the Chromium Next GEM Chip G (10X Genomics, Pleasanton, CA, USA). In total two pools of six samples each were subjected to partitioning into Gel bead-in-EMulsion on four Chromium Next GEM Chip G lanes followed by cDNA amplification and library preparation using the Chromium Single Cell 3' gene expression system (10X Genomics, Pleasanton, CA, USA). Final gene expression libraries were sequenced using a NextSeq 2000 System (Illumina, San Diego, CA, USA), with NextSeq 2000 P3 Reagents (100 Cycles) applying recommended sequencing conditions (read1: 28nt, i7 Index: 10nt, i5 index: 10nt, read2: 90nt). Sequencing output files were pre-processed using cell ranger (10X Genomics, Pleasanton, CA, USA). Downstream analyses were performed using software R, version 4.2 and 4.3. This included demultiplexing of cell hashing tags using the HTODemux function from package Seurat [5], removal of ambient RNA background using the decontX algorithm from the celda package, and multiplet exclusion using the R package Rscrublet, where multiple percentile variable numbers were tested to identify the optimal threshold for defining doublets. Quality control filtering including clustering was performed by removing cells with high mitochondrial content (≥20%), low RNA counts (≤300) and features (≤250), high RNA counts (≥60000) and features (≥8000), and clusters with a high proportion (>50%) of cells failing QC. In total, 8.7% of the cells were removed. The Seurat package was also used for normalization, variable feature selection, scaling, PCA, and UMAP dimensionality reduction and visualization. Differential expression analysis was conducted separately for endothelial and AT2 cells using EdgeR [6], with filtering for minimum proportions of cells expressing a gene (30%), minimum count (≥5) required for at least some samples, and minimum number of cells per sample after aggregating to pseudo-bulk profiles (≥8). For gene set enrichment analysis, we used R package clusterProfiler [7] requiring a minimum gene set size of 3, maximum gene set size of 800, and p-value cutoff of 0.05.

Multiplets were excluded after predicting them using Scrublet [8]. Cell-cell communication events between cell types were calculated using LIANA (LIgand-receptor ANalysis frAmework) [9, 10] as described previously [11] following single-cell best practice guide from Heumos *et al*. [12]. Murine genes were mapped to their human orthologs using the Alliance of Genome Resources database vs7.1.0 May 2024. Ligand-receptor pairs were filtered with specificity and magnitude ranks $\leq 0.05$, focusing on differentially-expressed genes from edgeR with a significance threshold of $p \leq 0.01$. Summed magnitudes of ligand-receptor interactions were calculated and compared between experimental conditions. The influence of ligands on potential target gene expression in receptor cell types was inferred using NicheNet [13], also mostly following single-cell best practice guide from Heumos *et al*. [12]. We used the Python packages LIANA+ [14], Scanpy [15]and AnnData [16] and R library nichenetr [13].

NicheNet analysis integrated results from AT2 cells, endothelial cells, neutrophils, and fibroblasts and considered both, up-regulated genes in wild-type, and in LIX knockout conditions. Predictions were filtered for regulatory potential ($\geq 0.035$), considering only differentially expressed genes as targets (EdgeR $p \leq 0.005$, absolute log2 fold change > 2).

**Spatial transcriptomics**

Whole lung tissue was fixed in 4% phosphate buffered formaldehyde solution (approx. 10% formalin solution) and paraffin embedded as described previously [17]. 5 µm thick sections of the left lungs were cut from formalin-fixed paraffin-embedded (FFPE) blocks, placed on Visium Spatial Gene Expression slides (10X Genomics, Pleasanton, CA, USA) and processed according to the manufacturer's instructions. H&E-stained tissue section images were generated using a Leica Aperio AT2 pathology scanner (Leica, Wetzlar, Germany) and captured at 40x magnification. Gene expression libraries were prepared according to manufacturer's instructions and sequenced using the NovaSeq X Plus System (Illumina, San Diego, CA, USA) and NovaSeq X Series 10B Reagent Kit applying recommended sequencing conditions. Sequencing output files were pre-processed using space ranger software (10X Genomics, Pleasanton, CA, USA) and downstream analyses were performed using software R, version 4.3 with the Seurat package [5]. Unsupervised clustering identified 11 distinct clusters which matched with the following histopathological areas: intact uninfected peripheral areas (cluster 0), intact infected peripheral areas (cluster 2), intermediate (cluster 3) and highly infiltrated areas (cluster 1) as well as edematous areas (cluster 6).

#### Data Availability

Single-cell RNA sequencing and Spatial transcriptomics data files are publicly available through Zenodo [redacted], the code used for data analysis is available at github.com, https://github.co m/GenStatLeipzig/Neutrophil-chemoattractant-CXCL5-contributes-to-loss-of-lung-barrier-integrity-in-acute-lung-injury.

#### Isolation of murine primary cells for cell culture

Naïve female and male C57BL/6J wild-type (WT) mice aged 8-16 weeks were used for cell isolation. Mice were anesthetized with ketamine and xylazine and then exsanguinated, as previously described [18]. Murine alveolar macrophages from bronchoalveolar lavage (BAL) were isolated and seeded onto 24-well cell culture plates as previously described [19]. Murine lung epithelial cells were isolated as previously described from dispase-digested lung tissue followed by negative magnetic beads separation and subsequent culture with epithelia-specific media for 5 days [20].

#### Cells and cell culture conditions

All cells were kept at 37°C in a 5% $CO_2$ atmosphere. Murine epithelial cell line - T7 cells (ECACC 07021402) were grown in DMEM media (Life Technologies, Carlsbad, CA, USA) supplemented with 10 % FCS (heat inactivated, CAPRICORN Scientific, Ebsdorfergrund, Germany), 0.2 mM L-Glutamine (Life Technologies, Carlsbad, CA, USA), 10 mM HEPES (Life Technologies, Carlsbad, CA, USA) and 1 % insulin-transferrin-selenium A supplements (Life Technologies, Carlsbad, CA, USA). Prior to infection and “cell stretch” cells were harvested and seeded in Opti-MEM (Life Technologies, Carlsbad, CA, USA). Human bronchial epithelial cell line BEAS-2B (ATCC, CRL-9609™) were cultured in Keratinocyte-SFM medium (Thermo Fisher Scientific, Waltham, MA, USA), containing 1 % penicillin/streptomycin (Life Technologies, Carlsbad, CA, USA). Prior to infection cells were harvested and seeded in Keratinocyte-SFM medium without penicillin/streptomycin. Human primary alveolar epithelial cell (HPAECs; H-6053, Cell Biologics, Chicago, IL, USA) were grown in complete human epithelial cell medium (H6621, Cell Biologics, Chicago, IL, USA) supplemented with 5% FCS and antibiotics. Prior to infection cells were washed twice with PBS and supplied with complete human epithelial cell medium without FCS and antibiotics.

#### Cell infection assays

All cells were kept at 37°C in a 5% $CO_2$ atmosphere. Primary cells and cell lines were infected with *S.pn.* serotype 2 (D39) obtained from early log phase liquid culture. Bacteria were grown in Todd Hewitt broth (BD Biosciences, Heidelberg, Germany) containing 0.5 % yeast extract (BD Biosciences, Heidelberg, Germany) and 10 % FCS. Single colonies of *S.pn.* were resuspended in culture media at indicated multiplicity of infection (MOI). Cell culture supernatants were harvested at indicated time points post-infection (p.i.) and stored at -80 °C until further analysis.

**Mechanical cell-stretch assays**

Mechanical stretch was applied to cells by Flexcell® FX-5000T™ or FX-6000T™ Tension Systems (FlexCell International, Burlington, NC, USA). Briefly, cells were seeded on six-well fibronectin or collagen IV coated BioFlex™ culture plates. Cyclic stretch was applied in a sinusoidal pattern with 18% amplitude at 0.25 Hz for the designated duration (stretch group). Cells cultured in the same plates but left non-stretched were used as controls (static group).
Murine T7 cells were stretched according to the following protocol: Cells were seeded on collagen IV coated BioFlex™ culture plates, grown for 24h without stretch and then either stretched for 24 h (18%, 0.25 Hz) or kept at static conditions.
HPAECs (HPAECs, H-6053, Cell Biologics, Chicago, IL, USA) were stretched according the following protocol: Cells were seeded on fibronectin-coated (100 µg/ml) BioFlex™ culture plates, grown for 24h without stretch and then either stretched for 24h (18%, 0.25 Hz) or kept at static conditions.

**Measurement of transepithelial electrical resistance (TEER)**

Human primary alveolar epithelial cells at passage 11-12 (HPAECs, H-6053, Cell Biologics, Chicago, IL, USA), or human pulmonary microvascular endothelial cells at passage 6-8 (HPMECs, Promocell, Heidelberg, Germany) were seeded at a density of 20,000 cells/well or 10,000 cells/well, respectively, on 96-well plates (96W10idf PET, Applied Biophysics, Troy, NY, USA) After 48-72h, TEER was calculated from electric cell impedance sensing (ECIS® Z-Theta Applied Biophysics, Troy, NY, USA) at 900 Hz for HPAECs or 4,000 Hz for HPMECs every 60 sec. Prior measurement media was replaced by Opti-MEM™ (Life Technologies, Carlsbad, CA, USA) for HPAECs or to basal endothelial media for HPMECs. After 1 h stabilization, 10 ng/ml TNF (PeproTech, Hamburg, Germany) was added for a preincubation time of 10min, followed by application of 100 ng/ml hCXCL5/ENA-78 (5-78aa, PeproTech, Hamburg, Germany) or hCXCL5/ENA-78 (8-78aa, PeproTech, Hamburg, Germany). Full-length hCXCL5 is 78 amino acids (aa) in length, but N-terminal proteolysis generates shorter variants with differential potency [21]. For our assays, we used a long-length hCXCL5(5-78aa) and a short-length hCXCL5(8-78aa) (Figure 6, E8). Cell barrier integrity was monitored continuously for 12h. Resistance was normalized for each well to the baseline before treatment.

**Immunofluorescence staining**

HPAECs and HPMECs were seeded on 8 well μ-slides (ibidi) and grown to confluence. Media was replaced by Opti-MEM™ (Life Technologies, Carlsbad, CA, USA) for HPAECs or by basal endothelial media for HPMECs. After 1h stabilization, 10 ng/ml TNF (PeproTech, Hamburg, Germany) was added for a pre-incubation time of 10 min, followed by application of 100 ng/ml hCXCL5/ENA-78 (5-78aa, PeproTech, Hamburg, Germany) or hCXCL5/ENA-78 (8-78aa, PeproTech, Hamburg, Germany) for 3h or 12h and fixed with 4% phosphate buffered formaldehyde solution. Fixed monolayers were permeabilized, quenched, blocked, and incubated with VE-cadherin (D87F2) XP® Rabbit mAb (1:400, #2500, Cell Signaling Technology, Danvers, MA, USA) and FITC-E-Cadherin (1:1000, 612131, BD Biosciences, Heidelberg, Germany) overnight at 4°C. After washing with Tris-buffered saline with 0.1% Tween® (TBS-T), cells were incubated with AF488-goat anti-rabbit secondary antibody (1:1000, A11008, Invitrogen, Waltham, MA, USA) and AF546-phalloidin (1:400, A22283, Thermo Fisher Scientific, Waltham, MA, USA) for 1h at room temperature. Nuclei were counterstained with DAPI at 1 μg/ml and imaged using a Scanning Confocal A1Rsi+ (60x Plan Apo objective, Nikon, Tokyo, Japan).

**Statistical analysis**

PRISM Graphpad software (version 9 and 10) was applied for statistical investigations. Survival analyses were performed using Kaplan-Meier curves and log-rank (Mantel-Cox) test. For parametric data, two groups were compared using unpaired *t* test, three or more groups by one-way ANOVA (Kruskal-Wallis test). For non-parametric data, two groups were compared using Mann Whitney test. Grouped analyses of parametric and non-parametric data were performed using two-way ANOVA/Tukey’s multiple comparison test. *P* values < 0.05 were considered statistically significant.

#### EXTENDED REFERENCES

#### SUPPLEMENTAL TABLES

**Suppl. Table E1: Metadata for patients with CAP and non-CAP controls in this study.** Values are given as the mean rounded to the next integer (if applicable). Enclosed in brackets is the range for the given data point. Day 0 refers to day of inclusion into study. Oxygenation index ($PaO_2/FiO_2$) n = 6 (except in days 0 and 6; n = 5).

| Variable | Cohort | Non-CAP |
|---|---|---|
| **N** | 6 | 5 |
| Female/Male | 5/1 | 3/2 |
| Age (years) | 54 (35-78) | 41 (27-71) |
| Height (cm) | 167 (160-170) | not obtained |
| Weight (kg) | 74 (55-105) | not obtained |
| Body mass index (kg/m$^2$) | 27 (20-41) | not obtained |
| Current smoker | 2 | 3 |
| Former smoker | 1 | 0 |
| Oxygenation index average over study time (mmHg) | Day 0: 219 (95-320) | not obtained |
| | Day 1: 239 (156-308) | |
| | Day 2: 277 (220-367) | |
| | Day 3: 218 (119-340) | |
| | Day 4: 196 (92-277) | |
| | Day 5: 219 (149-274) | |
| | Day 6: 267 (186-329) | |
| Oxygenation index average at day of lavage (mmHg) | 225 (95-320) | not obtained |
| **Comorbidity** | | |
| COPD | 1 | not obtained |
| Diabetes | 1 | not obtained |
| **Scores** | | |
| CURB-65 | 3 (2-4) | not applicable |
| ATS minor criteria | 5 (4-6) | not applicable |
| PSI class | 3 (1-5) | not applicable |
| SOFA score | 17 (15-20) | not applicable |

#### SUPPLEMENTAL FIGURES

##### Figure E1

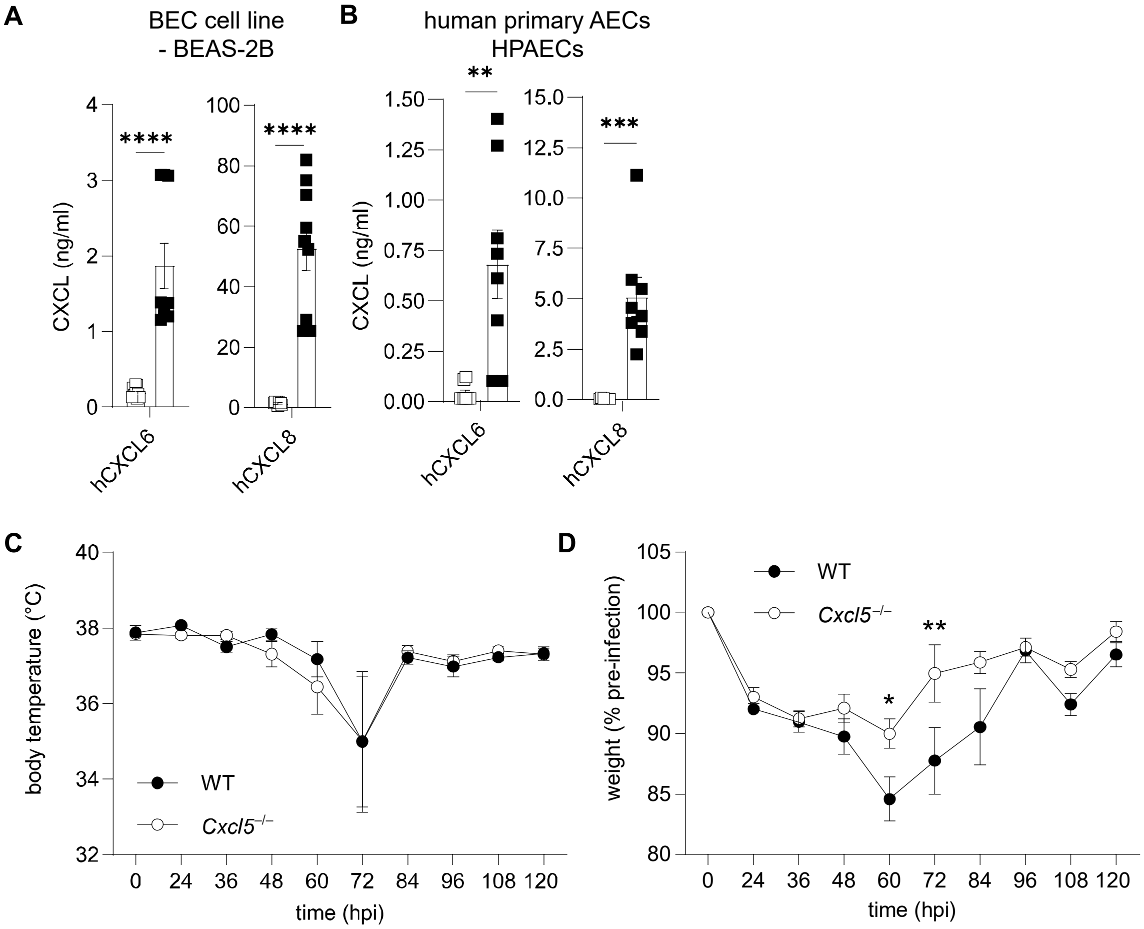


**Figure E1. Human CXCL chemokine production by BEAS-2B and HPAECs and body temperature and body weight curves of mice with pneumococcal pneumonia.** Human CXCL6 and CXCL8 chemokine concentrations measured in cell culture supernatants 24 hours post infection (hpi) by ELISA in (**A**) BEAS-2B BEC line following *S.pn.* infection (MOI 0.1, n = 9, pooled from 3 experiments) and (**B**) HPAECs following *S.pn.* infection (MOI 1, n = 8, pooled from 4 experiments). Mean ± SEM. Mann Whitney test, unpaired. ** *P* < 0.01, *** *P* < 0.001 and **** *P* < 0.0001. BEC, bronchial epithelial cells; MOI, multiplicity of infection. (**C**, **D**) WT (n = 12) and *Cxcl5*$^{-/-}$ mice (n = 14) mice were intranasally infected with 5 x $10^6$ CFU *S.pn.* ST2 and monitored up to 120 hpi. Data are pooled from two experiments. (**C**) Body temperature and (**D**) body weight (% of pre-infection weight) of mice monitored in 12h intervals. Mixed-effects model (REML). Šídák's multiple comparisons test. * *P* < 0.05 and ** *P* < 0.01.

### Figure E2

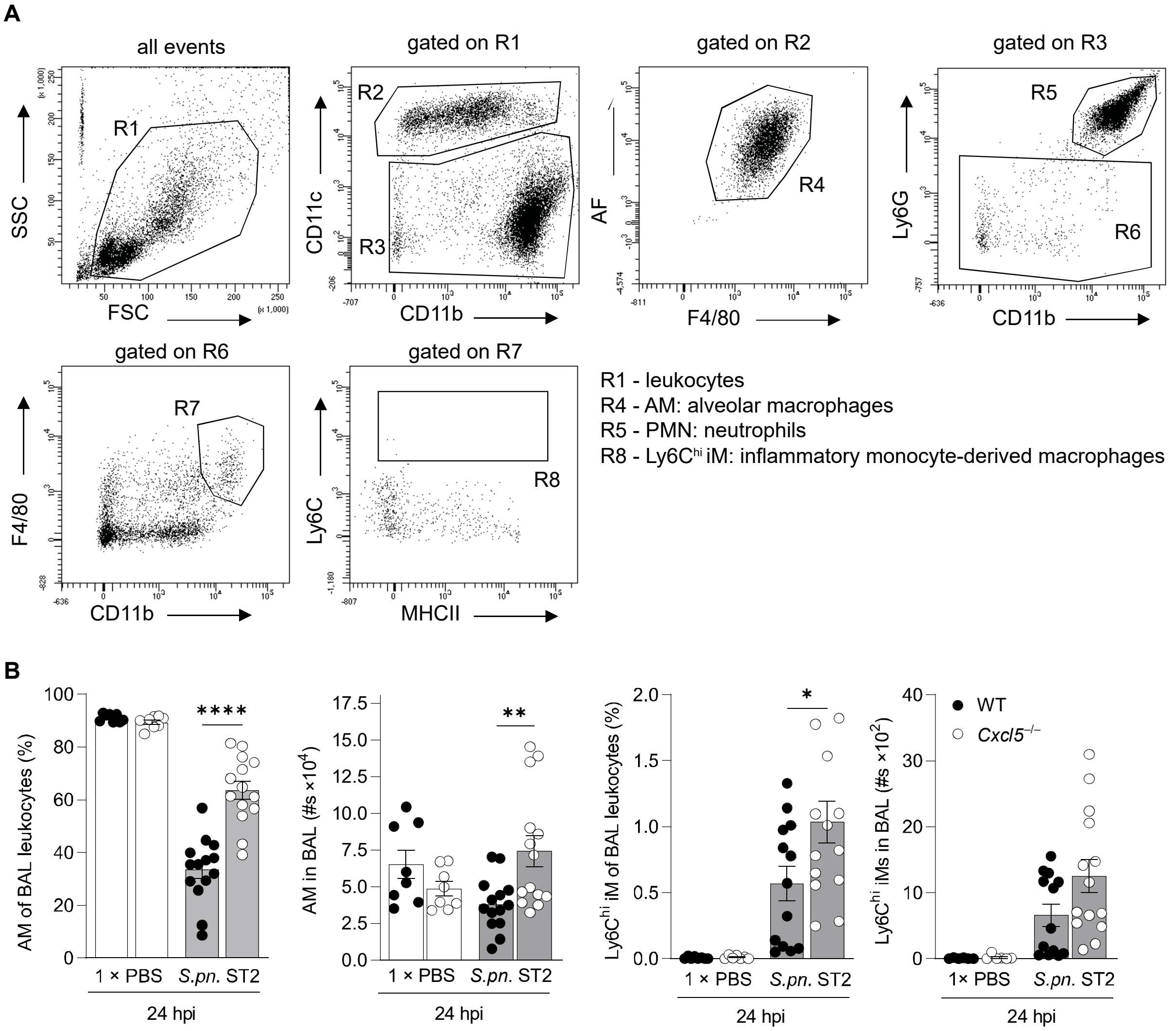


**Figure E2. Gating strategy and alveolar leukocytes recruitment.** (**A**) Gating strategy to discriminate alveolar macrophages (AM), neutrophils (PMN) and inflammatory monocyte-derived macrophages (Ly6C$^{hi}$ iM), 24 hpi time point analysis of *S.pn.* ST2 infected WT (n = 14), *Cxcl5*$^{-/-}$ (n = 14) and PBS ctr WT (n = 8) and *Cxcl5*$^{-/-}$ (n = 8) mice, data pooled from 2 – 3 experiments, mean ± SEM. (**B**) Frequencies and total numbers of BAL AMs and Ly6C$^{hi}$ iMs in *S.pn.*- or PBS-infected WT and *Cxcl5*$^{-/-}$ mice. Mean ± SEM. One-way ANOVA, Šídák's multiple comparisons test, WT vs *Cxcl5*$^{-/-}$ mice. (**B**), Ly6C$^{hi}$ iMs % and #s outliers identified by ROUT method were removed (WT-*S.pn.*$_{\%}$= 4.2% and WT-*S.pn.*$_{\#s}$= 138.9 x $10^2$ cells; *Cxcl5*$^{-/-}$-PBS$_{\%}$= 0.18% and *Cxcl5*$^{-/-}$-PBS$_{\#s}$= 0.986 x $10^2$ cells and WT-PBS$_{\#s}$= 0.171 x $10^2$ cells, 0.138 x $10^2$ cells). One-way ANOVA. Šídák's multiple comparisons test. * $P < 0.05$, ** $P < 0.01$ and **** $P < 0.0001$. BAL, Bronchoalveolar lavage; hpi, hours post infection; WT, wild type; *S.pn.*, *Streptococcus pneumoniae;* AMs, alveolar macrophages; Ly6C$^{hi}$ iMs, = inflammatory monocyte-derived macrophages.

#### Figure E3

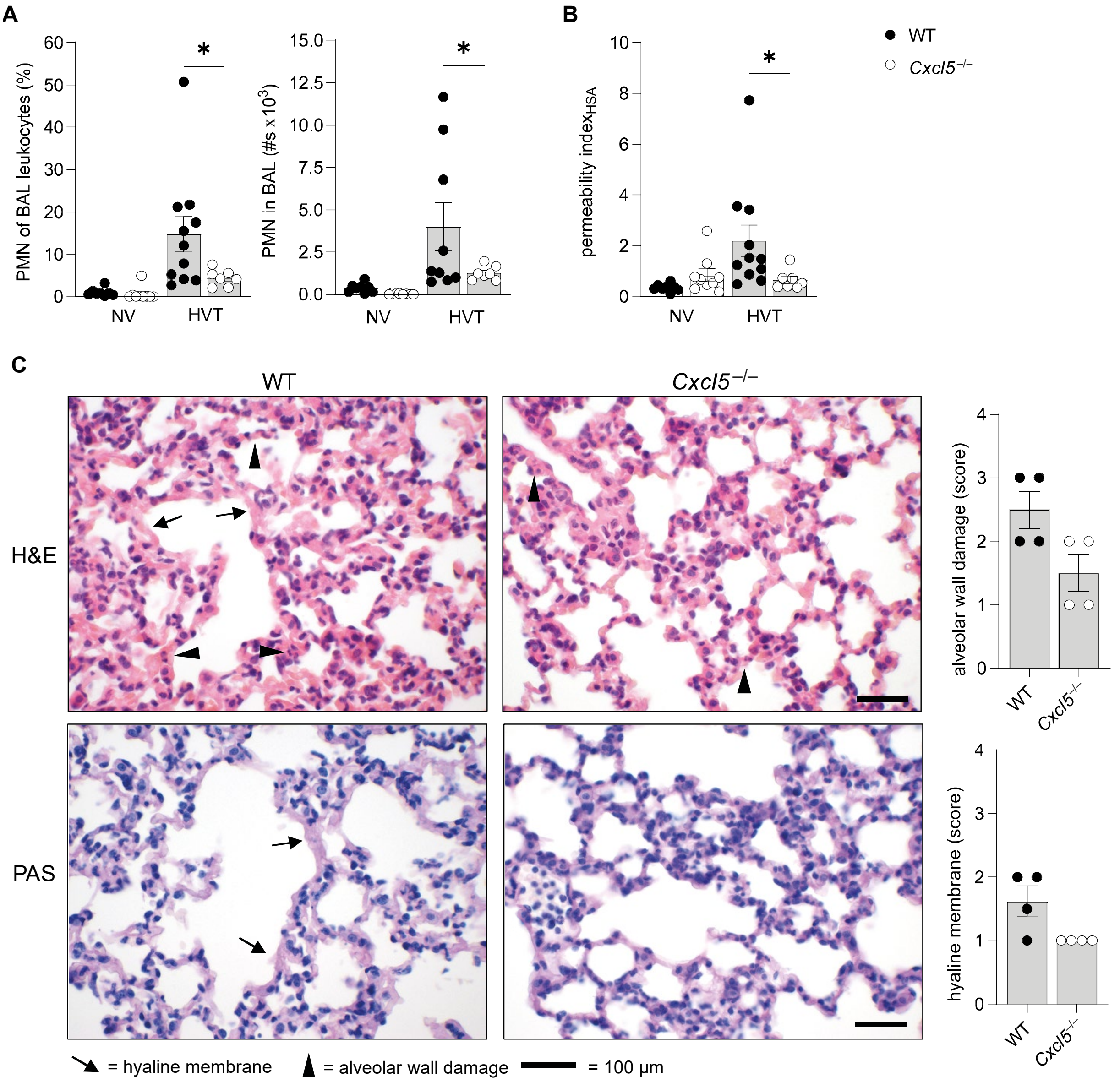


**Figure E3. Absence of CXCL5 results in reduced neutrophil recruitment and preservation of alveolar-capillary barrier integrity in murine ventilator-induced lung injury model.** (**A**, **B**) Alveolar neutrophil recruitment (**A**) and barrier permeability index (**B**) in WT (n = 11) and *Cxcl5*$^{-/-}$ (n = 7) mice following 4h of high tidal volume (34 ml/kg body weight) mechanical (HVT) or 5-10 min low tidal volume (9 ml/kg body weight) control ventilation (NV) (n = 8 and 9). (**A**) Frequencies and total numbers of BAL PMN following HVT or NV. PMN #s outliers identified by ROUT method were removed (WT-HVT # = 39.3 x $10^3$ cells and Cxcl5$^{-/-}$NV = 1.17 x $10^3$ cells, for 1 WT-HVT only frequencies were measured. (**B**) Alveolar permeability index of HSA following HVT or NV. (**C**) H&E and PAS staining of WT (n = 4) and *Cxcl5*$^{-/-}$ (n = 4) murine lungs depicting alveolar wall damage (indicated by arrow heads) and hyaline membrane (arrows) following HVT. (**A – B**) Data displayed as mean ± SEM, (**C – D**) graphs displayed as median with interquartile range. (**A – B**) One-way ANOVA, Šídák's multiple comparisons test, HVT WT vs *Cxcl5*$^{-/-}$ mice. Mean ± SEM. * $P < 0.05$, ** $P < 0.01$. (**C**) Scale bare indicates 100 μm. Error bars indicate mean ± SEM.

#### Figure E4

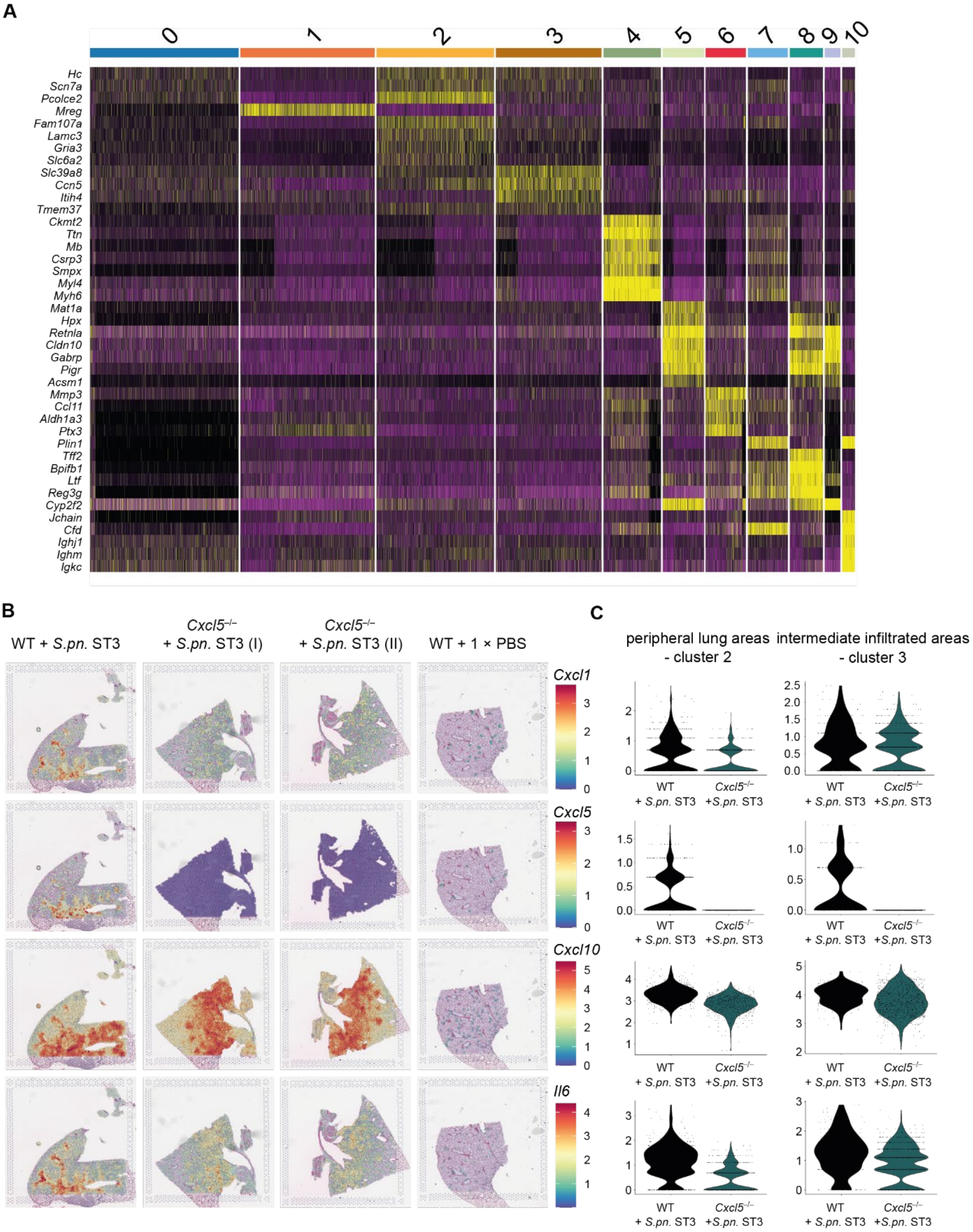


**Figure E4. Spatially resolved transcriptomic profiling of lung tissue sections.** (**A**) Heatmap of cluster defining genes (top marker genes for every cluster). Clusters matched with the following histopathological areas: intact uninfected peripheral areas (cluster 0), intact infected peripheral areas (cluster 2), intermediate (cluster 3) and highly infiltrated areas (cluster 1) as well as edematous areas (cluster 6). (**B**) "SpatialFeaturePlots" displaying the expression of indicated genes: "*Cxcl1*", "*Cxcl5*", "*Cxcl10*" and "*Il6*" in lung tissue sections of *S.pn.* ST3 infected WT (n = 1), *S.pn.* ST3 infected *Cxcl5*$^{-/-}$ (n = 1, two sections) and PBS ctr WT (n = 1) mice at 24 hpi. (**C**) Violin plots displaying gene expression of "*Cxcl1*", "*Cxcl5*", "*Cxcl10*" and "*Il6*" in indicated clusters.

#### Figure E5

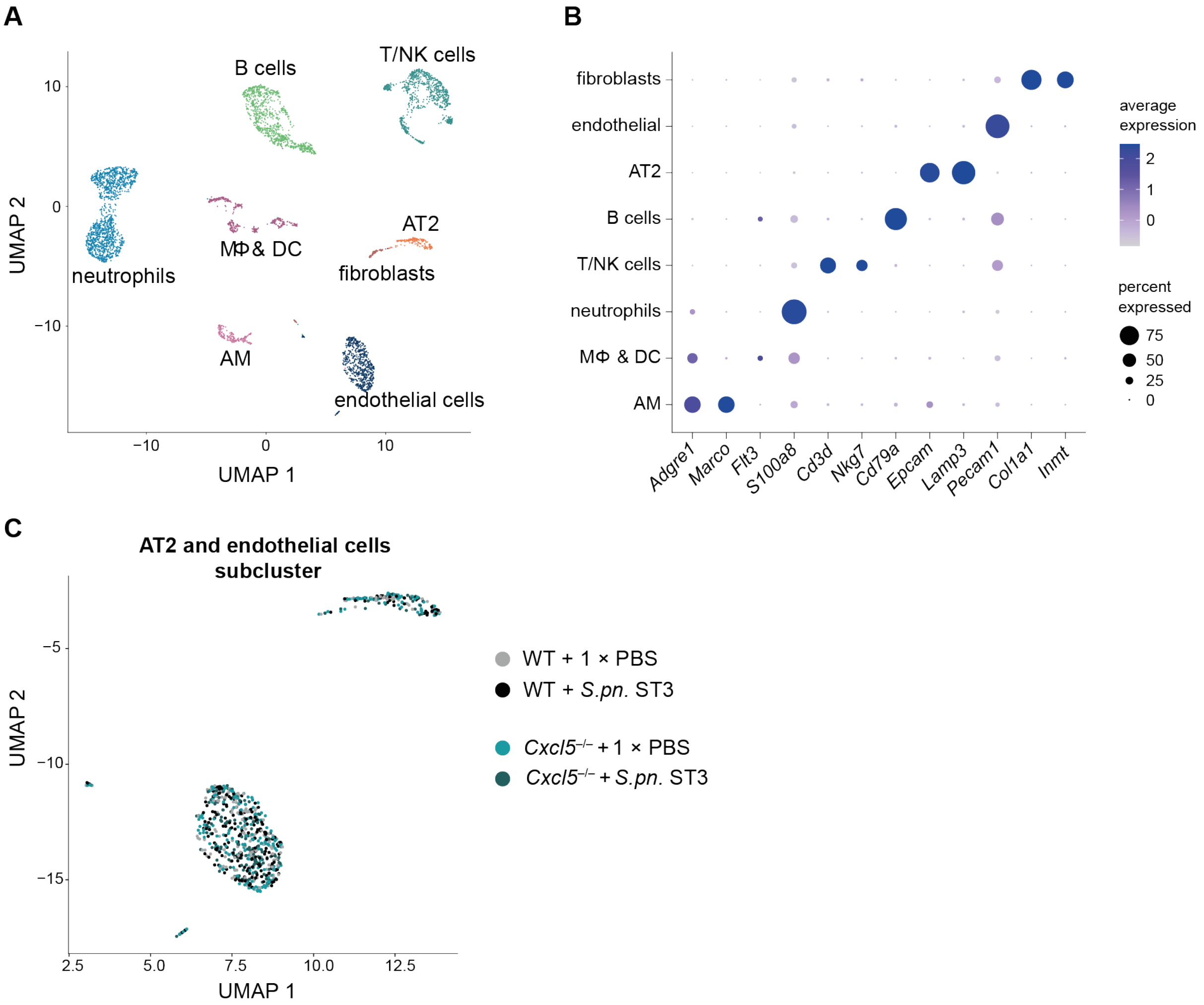


**Figure E5. Single-cell gene expression analysis in lungs.** (**A**) Uniform manifold approximation and projection (UMAP) plot of sequenced lung cells isolated from *S.pn.* ST3-infected or PBS ctr WT (n = 3) and *Cxcl5*$^{-/-}$ (n = 3) mice 24 hpi and following data integration. (**B**) Marker genes used for annotation of unsupervised lung cell clusters. (**C**) UMAP plot of alveolar epithelial cells type 2 (AT2) and endothelial cell subset colored by experimental groups. MΦ, macrophages; DC, dendritic cells; AM, alveolar macrophages; NK, natural killer cells; AT2, alveolar epithelial cells type 2.

### Figure E6

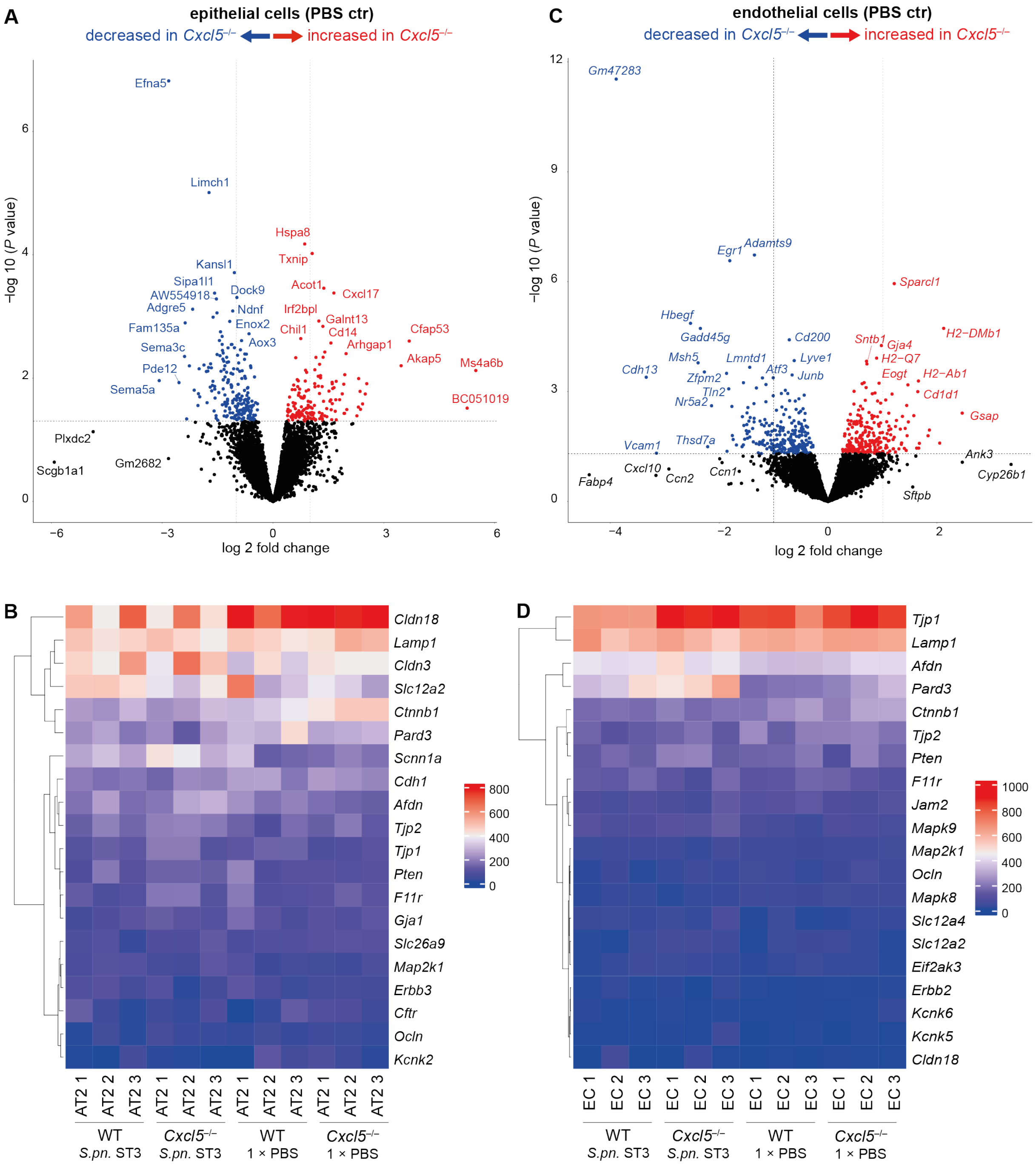


**Figure E6. Transcriptomic profile of lung epithelial and endothelial cells.** (**A**) Volcano plot displaying differentially expressed genes of AT2 cells between PBS ctr WT (n = 3) and *Cxcl5*$^{-/-}$ (n = 3) mice 24 hpi. (**B**) Heatmap displaying genes linked to barrier functions in AT2 cells of *S.pn.* ST3-infected or PBS ctr WT (n = 3) and *Cxcl5*$^{-/-}$ (n = 3) mice 24 hpi. (**C**) Volcano plot displaying differentially expressed genes of endothelial cells between PBS ctr WT (n = 3) and *Cxcl5*$^{-/-}$ (n = 3) mice 24 hpi. (**D**) Heatmap displaying genes linked to barrier functions in endothelial cells of *S.pn.* ST3-infected or PBS ctr WT (n = 3) and *Cxcl5*$^{-/-}$ (n = 3) mice 24 hpi. (**A**, **C**) Colored dots indicate differential expressed genes with *P* values below 0.05. Dashed lines indicate *P* value of 0.05 (horizontal) or log2-fold changes of -1 and 1 (vertical). (**D**) Analogous to (**B**) for endothelial cells. Differential gene expression analysis were performed using the edgeR framework [6].

#### Figure E7

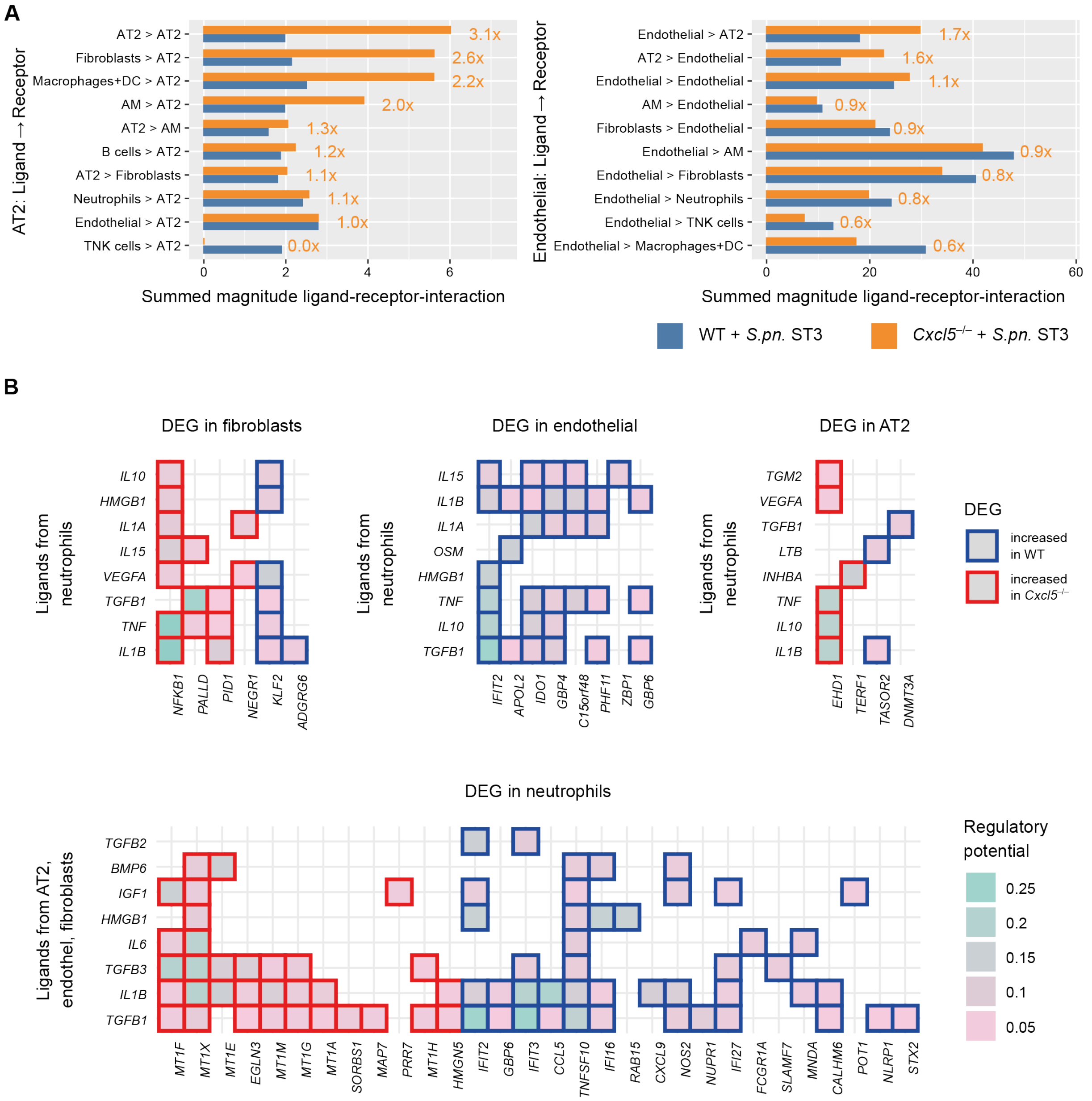


**Figure E7. Cell-cell communication analysis of predicted ligand-receptor interactions and ligand-target-gene interactions.** (**A**) Shown is the sum of interaction magnitudes amongst lung single cells when acting as either target or source, respectively, in *S.pn.* ST3-infected WT (n = 3) and *Cxcl5*–/– (n = 3) mice at 24 hpi fpr the top-10 ligand-receptor pairs ordered by summed magnitude. Orange numbers report the ratio between *Cxcl5*–/– and WT. (**B**) Prediction of top-differentially expressed genes (DEGs) ordered by regulatory potential induced in target cells by released ligands from specified cell types. [9, 10] and NicheNet [13] framework. Depicted genes are predicted human orthologs of murine genes.

#### Figure E8

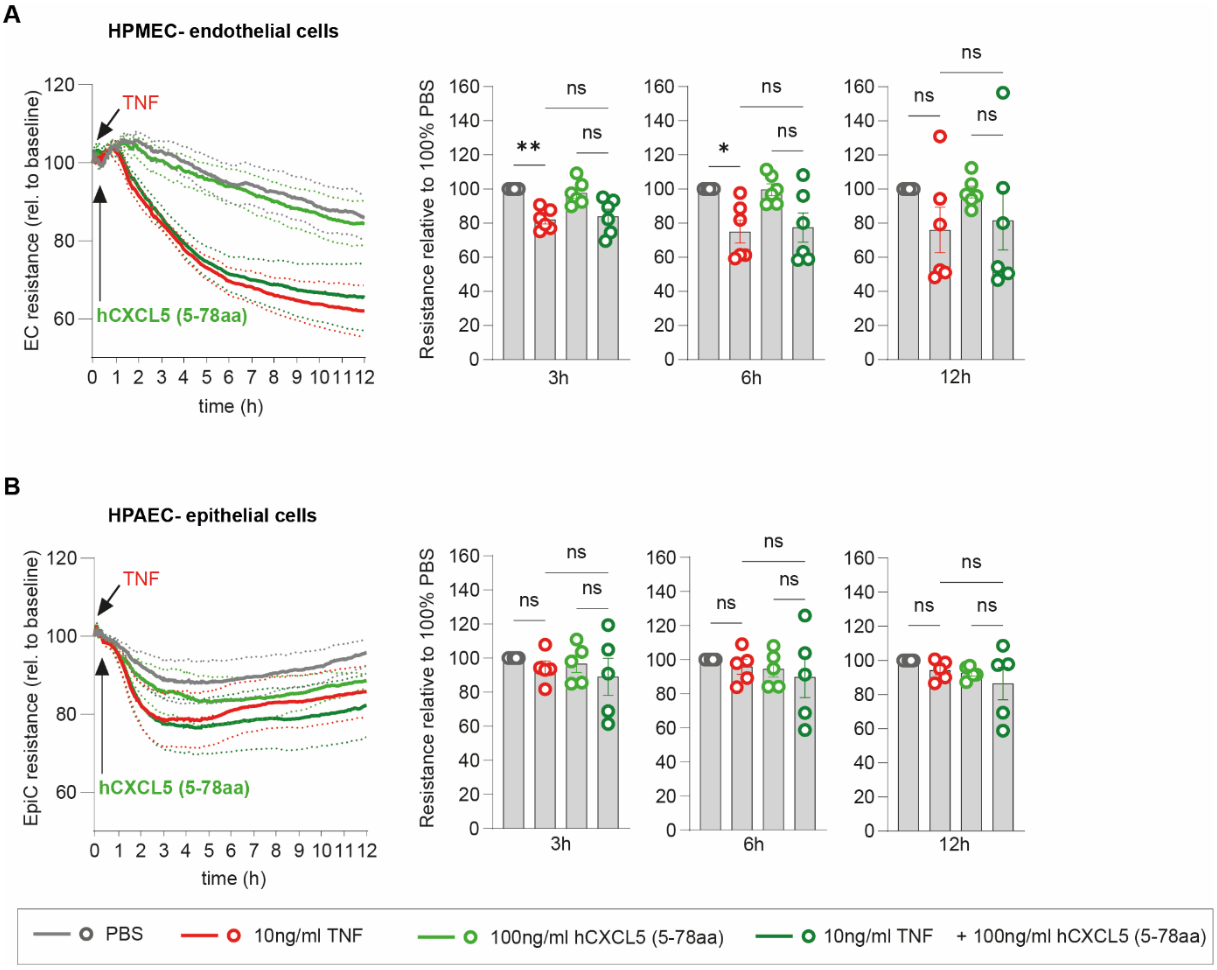


**Figure E8. Human CXCL5(5-78aa) stimulation of TNF-primed human primary alveolar epithelial cells and pulmonary microvascular endothelial cells**. (**A**) Human pulmonary microvascular endothelial cells (HPMEC) and (**B**) Human primary alveolar epithelial cells (HPAEC) were primed with 10 ng/ml TNF, followed by stimulation with 100 ng/ml hCXCL5 (5-78aa). Data are presented as mean ± SEM; for curve analysis, two-way ANOVA, Dunnett's multiple comparisons test, or for comparison of normalized resistance values at the designated timepoints, Kruskal Wallis-test was performed. *P* values indicated in Figure.
